# Opposing diet, microbiome and metabolite mechanisms regulate inflammatory bowel disease in a genetically susceptible host

**DOI:** 10.1101/2022.04.03.486886

**Authors:** Gabriel Vasconcelos Pereira, Marie Boudaud, Mathis Wolter, Celeste Alexander, Alessandro De Sciscio, Erica. T. Grant, Bruno Caetano Trindade, Nicholas A. Pudlo, Shaleni Singh, Austin Campbell, Mengrou Shan, Li Zhang, Qinnan Yang, Stéphanie Willieme, Kwi Kim, Trisha Denike-Duval, Jaime Fuentes, André Bleich, Thomas M. Schmidt, Lucy Kennedy, Costas A. Lyssiotis, Grace Y. Chen, Kathryn A. Eaton, Mahesh S. Desai, Eric C. Martens

## Abstract

Inflammatory bowel diseases (IBDs) are chronic conditions characterized by periods of spontaneous intestinal inflammation and are increasing in industrialized populations. Combined with host genetics, diet and gut bacteria are thought to contribute prominently to IBDs, but mechanisms are still emerging. In mice lacking the IBD-associated cytokine, Interleukin-10, we show that low dietary fiber promotes bacterial erosion of colonic mucus, leading to lethal colitis. A fiber-free exclusive enteral nutrition diet also induces mucus erosion but inhibits inflammation by simultaneously increasing an anti-inflammatory bacterial metabolite, isobutyrate. Diet-induced inflammation is driven by Th1 immune responses, which increase in the presence of mucin-degrading bacteria and are preceded by expansion of natural killer cells and altered immunoglobulin-A coating of some bacteria. Inflammation occurs first in intestinal regions with thinner mucus. Our work underscores the importance of focusing on microbial functions—not taxa—contributing to IBDs and some diet-mediated functions block those that promote disease.

## Introduction

Inflammatory bowel diseases (IBDs) are characterized by periods of spontaneous inflammation in the gastrointestinal tract and occur in people with underlying genetic variations that cause inappropriate immune responses to intestinal antigens, especially ordinarily non-harmful symbiotic gut microbes^1^. Despite identification of hundreds of IBD-associated gene polymorphisms^2^, precise mechanisms through which IBDs develop have not been determined. Even when predisposing genetics exist, IBDs do not always occur, suggesting that additional important triggers beyond host genetics and gut microbes are required^3^.

The incidence of IBDs is increasing in some industrializing countries^4^ and in immigrant populations that move to industrialized countries^5^. Dietary changes associated with industrialization (decreased fiber, increased processed foods, emulsifiers and sugars) are emerging as potential “triggers” that enhance susceptibility^6–9^ and some underlying mechanisms are beginning to emerge^7,9^. The physiology of the microbes inhabiting the gut is continuously influenced by diet, especially fiber polysaccharides that elude digestion in the upper gastrointestinal tract and arrive in the colon providing nutrients for microbes^10–12^. Several studies using mice fed low fiber, high sugar or other “Westernized” diets have shown alterations in the microbiome that correlate with reduced integrity of the mucus barrier, including increased activity of mucin-degrading bacteria and reduced mucus thickness^9,13, ,14,15,16^ and increased mucus penetrability^17^.

We investigated the contributions of dietary fiber and mucin-degrading gut bacteria to the development of inflammation in mice lacking the IBD-associated cytokine Interleukin-10 (IL-10). In humans, loss of IL-10 or either of its receptor subunits results in early onset IBD in infants and children^18^. Conventional *Il10^-/-^* mice develop spontaneous inflammation that is variable between mouse colonies and worsened by the presence of pathobionts like *Enterococcus faecalis* or *Helicobacter* spp.^19^, while deriving mice as germfree reduces or eliminates inflammation^20^. Thus, gut microbes are required for inflammation in *Il10^-/-^* mice, however, the mechanisms mediating disease progression in the presence of commensal bacteria lacking known pathogenic qualities remain unknown.

We found that a combination of IL-10 loss, gut microbiota containing mucin-degrading bacteria and a low-fiber diet, which promotes bacterial mucus erosion, are all required to elicit severe intestinal inflammation. We also observed that the clinically validated diet therapy exclusive enteral nutrition (EEN) partially inhibits inflammation—despite lacking dietary fiber and promoting mucus erosion—in part by promoting production of the bacterial metabolite isobutyrate. Our work highlights the idea that microbial *functions* should be the focus of positive and negative impacts in the development of IBDs. Some of these functions, like mucus degradation, can be genetically complex and difficult to predict from taxonomic or metagenomic data, necessitating the use of simplified systems in which individual functions contributing to IBDs can be investigated at mechanistic levels prior to research in humans.

## Results

### A combination of host genetic susceptibility, low dietary fiber and intestinal bacteria exacerbate colitis in Il10^-/-^ mice

We previously determined that colonizing wild-type (WT) germfree mice with a synthetic microbiota containing 14 species (SM14) of sequenced and metabolically characterized human gut bacteria results in erosion of colonic mucus and increased *Citrobacter rodentium* susceptibility in mice fed a fiber-free (FF) diet^13^. Fiber-deprived WT mice do not develop spontaneous, histologically evident inflammation. Notably, mice colonized with the same SM14 and fed a fiber rich diet do not experience mucus erosion because metabolism of fiber by gut bacteria decreases the abundance and activity of mucin-degrading species^13^. To determine if diet- and microbiota-driven mucus erosion—and the correspondingly increased proximity of bacteria to host tissue—promotes disease in mice that are genetically susceptible to IBD, we introduced the SM14 into germfree *Il10^-/-^* mice, choosing the C57BL/6J background that has been reported to be *most* resistant to inflammation^19^. We colonized adult 7–10-week-old *Il10^-/-^* mice fed a fiber-rich (FR) diet, and switched a subset of the mice to the FF diet after 14 days of colonization, monitoring body weight up to 60d (**Figure 1A**). Mice kept on the FR diet (n=14) maintained or gained weight. In contrast, mice switched to the FF diet began losing weight 1–2 weeks after the diet switch and experienced 89% lethality by 60d (n=27). Histology revealed neutrophil infiltration, mucosal erosion, ulceration and edema in the ceca of FF-fed mice, but not those fed FR (**Figure 1B**). These markers were lower in the ileum and colon (**Figure 1C**). Cecal inflammation did not develop in SM14-colonized WT mice fed either diet (**Figure 1D**). Measurements of neutrophil-derived Lipocalin-2 (LCN2) in the cecal lumen provided additional data supporting severe inflammation in SM14-colonized *Il10^-/-^* mice fed the FF diet (**Figure 1E**). Experiments in which individual variables for diet (FR, FF), colonization (SM14, germfree) and host genotype (WT, *Il10^-/-^*) were individually manipulated support the conclusion that severe inflammation and weight loss only develop in the context of three conditions: IL-10 deficiency, SM14 colonization and FF diet (**Figures 1D,E, S1A–D)**.

**Figure 1.**
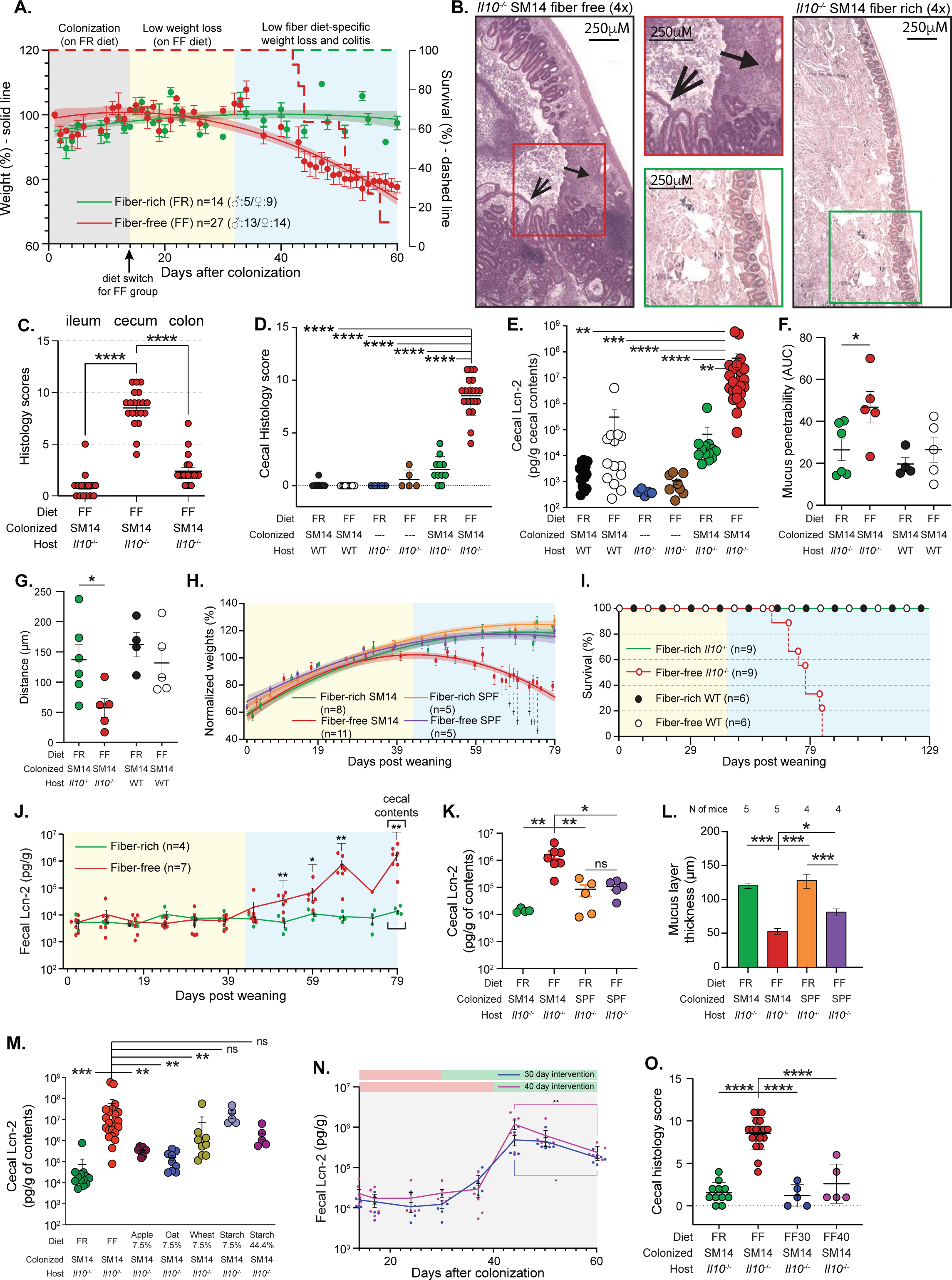
Low fiber driven inflammation in *Il10^-/-^* mice. **A.** Weights of adult *Il10^-/-^*mice colonized with the SM14 and maintained on a fiber rich (FR) diet or switched to a fiber free (FF) diet at 14d. Curves represent polynomial (quadratic) equations fit to all of the weights gathered at various days. Weights were measured more frequently after 40d due to declining health in the FF group. Animals that were euthanized were excluded from the curve at later points. Two FF-fed animals from an early experiment were found dead and were not included in subsequent analyses. Right axis shows survival over time. **B.** Representative cecal histology of FF (left) and FR (right) fed SM14-colonized *Il10^-/-^* mice with higher magnification insets in the middle. Block arrows points to a large ulcer, line arrow to an area of edema. **C.** Quantitative, blinded histological scoring of ileal, cecal, and colonic tissue from colonized *Il10^-/-^*mice fed the FF diet (n=15–20, one-way ANOVA and post hoc test with Holm-Šídák’s multiple comparison test) **D.** Histological scoring of cecal tissue taken from colonized *Il10^-/-^* mice fed the FR and FF diets, along with additional treatments to manipulate diet, colonization and host genotype variables (n=5–20). **E.** Cecal Lipocalin-2 (LCN2) measurements in the same treatments shown in D (n=5– 23). **F.** Mucus penetrability by 1 um-sized beads in the distal colon of *Il10^-/-^* and WT mice fed the FR and FF diets. **G.** Distance of 1 um-sized beads from the epithelium in the same mice in F. **H.** Weights of *Il10^-/-^* mice born to either SM14 or SPF parents and weaned to FR or FF. By 79d post weaning, only the SM14/FF group (red) displays weight loss. Curves are fit to weight data as in A. **I.** Survival curves for 4 separate groups of WT C57bl/6 or *Il10^-/-^* mice colonized by parental transfer of SM14 bacteria at birth. **J.** Fecal LCN2 measurements over time in SM14 colonized *Il10^-/-^* pups weaned to FR and FF. **K.** Endpoint cecal LCN2 in SM14 and SPF colonized mice (n=4–7). **L.** Mucus thickness measurements in SM14 and SPF colonized mice. Sample size indicated below each treatment group (n=4–5). **M.** Cecal LCN2 in mice fed versions of the FF diet with glucose replaced by dietary fiber from apple, wheat, oat or soluble starch. Concentrations of each supplement are noted below the ingredient (n=5–23). **N.** Fecal LCN2 measurements over time in SM14 colonized adult mice shifted to FF at 14d post colonization and then shifted back to FR at either 30d or 40d. For each time point/treatment the mean is shown along with individual points and error bars represent S.E.M. **O.** Endpoint histology of the cecal tissue from mice shown in N. compared to mice maintained on FR or FF (n=5–20). Experiments in panels F. and G. were done at the University of Luxembourg; all others were done at the University of Michigan. In panels C, D, E, J, K, L, M, bold horizontal bar represents the mean and lighter error bars the S.E.M. *P* values: * ≤ 0.05; ** ≤ 0.01; ** ≤ 0.001; *** ≤ 0.001; **** ≤ 0.0001; ns = not significant. In panels D, E, K, L, M, N, O, two-way ANOVA and post hoc test with Original FDR method of Benjamini and Hochberg was used for statistics.

Switching SM14-colonized *Il10^-/-^* mice to the FF diet rapidly induced a shift in bacterial composition characterized by decreased fiber-degraders (*B. ovatus* and *E. rectale*) and increased abundance of two of the four known mucin-degraders, *A. muciniphila* and *B. caccae* (**Figure S1E**). A similar trend was observed in wild-type mice fed the FF diet, albeit with significantly higher levels of *Escherichia coli* and *Bacteroides thetaiotaomicron* in *Il10^-/-^* mice (**Figure S1F,G**). As expected, feeding the FF diet to *Il10^-/-^* mice resulted in reduced mucus thickness (**Figure S1H–K**). We hypothesize that this erosion of protective mucus is critical for IBD development by increasing bacterial contact with the epithelium and the dysregulated *Il10^-/-^*immune system. Supporting this idea, using bacteria-sized florescent beads in an *ex situ* mucus penetrability assay^21^ we determined that FF-fed *Il10^-/-^* mice have higher colonic mucus penetrability and closer proximity of 1µm-sized beads to the host epithelium compared to their FR-fed counterparts (**Figure 1F,G, S1L,M**). This increase in mucus penetrability was not as pronounced in wild-type mice with SM14 (**Figures 1F,G)**.

Human disease associated with IL-10 dysfunction often presents as early or very early onset IBD in children^18^. With this in mind, we modified our model to allow natural microbiota transfer in the neonatal period, which also enables direct comparison to conventional specific pathogen free (SPF) mice that are also colonized at birth. We colonized germfree *Il10^-/-^* adult parents fed the FR diet, allowing their pups to be exposed to the maternal SM14 beginning at birth. In pre-weaned pups, the SM14 took on a similar composition as adult FF fed mice (**Figure S1N**) due to the lack of fiber and the fact that milk oligosaccharides share linkages with mucin *O*-glycans^22^. Pups weaned to the FF diet began losing weight after ∼39 days post weaning (dpw) (**Figure 1H**) and experienced 100% mortality by 84 dpw (**Figure 1I**). In most experiments, we harvested FR- and FF-fed groups at 79 dpw (100d old in University of Michigan facility), in which case mortality among the FF mice was ∼82% and all mice weaned to the FR diet survived (**Figure S1O**). A separate group of FR-fed mice showed 100% survival when maintained for 129 dpw (150d total) as did wild-type mice fed either diet (**Figure 1I**). Weight loss in pups weaned to the FF diet corresponded with increasing fecal LCN2 that initiated around the same time (∼39 dpw) of declining weight (**Figure 1J**).

Interestingly, *Il10^-/-^* mice with a conventional SPF microbiota fed the FF diet did not lose weight as severely as those colonized with SM14 (**Figure 1H**). Compared to SM14-colonized mice fed the FF diet, SPF mice showed lower cecal LCN2 at 79 dpw (**Figure 1K**), and less histological damage (**Figure S1P**), revealing that the more complex, murine microbiota does not promote the same level of inflammation. However, SPF mice fed the FF diet *did* exhibit reduced mucus thickness, albeit significantly less than SM14-colonized mice (**Figure 1L**), a point addressed in more detail below.

### Restoration of dietary fiber inhibits inflammation

The FR and FF diets differ in several aspects of their composition beyond fiber (**Table S1**). To more directly test fiber’s contribution, we created versions of the FF diet in which 7.5% of the glucose it contains was replaced with an equivalent amount of food grade, pure fiber from oat, wheat or apple. High sugar has been shown to promote inflammation, including in *Il10^-/-^* mice, and this effect is reduced by replacing some sugar with starch that is digested by the host in the small intestine^9^. As a control for reducing the sugar in our fiber-supplemented diets, we created a diet containing 7.5% highly digestible starch, exchanging free glucose for a glucose polymer available to the host via upper GI digestion. The presence of 7.5% fiber from either of the three sources, but not starch, significantly reduced LCN2 (**Figure 1M**) as well as weight loss and histopathology (**Figure S2A,B**). Even when all of the glucose (44%) was replaced with digestible starch, adult mice colonized with the SM14 developed disease (**Figures 1M, S2A,B**). This high starch diet more closely approximates a fiber poor human diet, which would be expected to be rich in starchy foods, protein and fat, and still promotes inflammation, implying that direct free sugar availability to the gut microbiota is not a major component of disease development in this model. *Bacteroides ovatus*, a proficient degrader of the polysaccharides expected to be present in the supplemental fibers^23^, was one of the major responders, increasing between 2–3 fold in relative abundance and this increase occurred at the expense of mucin-degrading *Akkermansia muciniphila* and *Bacteroides caccae* (**Figure S2C–G**).

To determine if restoring dietary fiber to mice that had already been fed the disease-promoting FF diet is capable of blocking inflammation, we returned colonized adult mice to the FR diet at either 30d or 40d (16 or 26 days after being switched to FF). Both groups of mice that were returned to FR exhibited no lethal weight loss (**Figure S2H**) and LCN2 levels and histology at 60d were lower than mice maintained on the FF diet (**Figure 1N, O**). Interestingly, temporal fecal LCN2 measurements showed that both groups experienced a peak of inflammation after fiber had been restored, which then began declining, indicating that low fiber induces disruption to the host**–**microbe homeostasis in which inflammation is a lagging effect that can eventually be reset (**Figure 1N**).

### The status of the mucus layer is a critical determinant in developing inflammation

To more directly evaluate the disease-promoting role of mucin-degrading bacteria, we colonized adult *Il10^-/-^* mice with a simpler synthetic microbiota containing only the 10 species (“SM10”) previously shown to be unable to grow on mucin oligosaccharides^13^. SM10-colonized mice exhibited 100% survival on the FF diet until 60d post colonization (n=7), significantly lower cecal LCN2 at 60d (**Figure 2A**) and reduced histological damage (**Figure 2B**). Eliminating the mucin-degrading bacteria also decreased mucus thinning (**Figures 2C, S2I**). Notably, SM10 colonized mice still exhibited some inflammation (**Figure 2B**). Consistent with the idea that the absence of mucus-degrading bacteria slows, but does not fully block, inflammatory activation of the dysregulated *Il10*^-/-^ immune system, SM10-colonized mice that were switched to the FF diet for 136d (150d total colonization) exhibited significantly better survival (50%) compared to FF-fed SM14 mice (**Figure 2D**).

**Figure 2.**
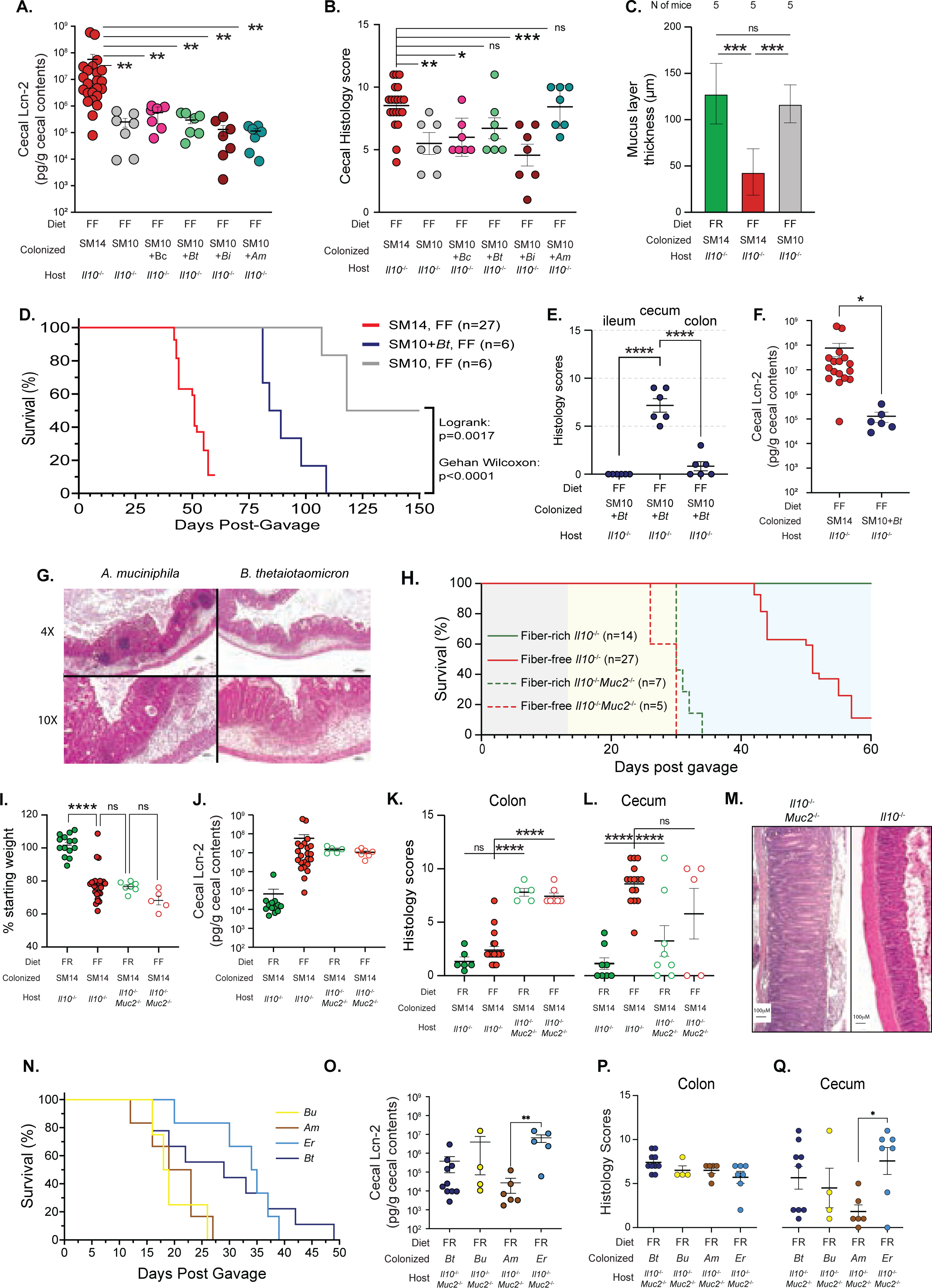
Mucus integrity is central to diet-induced inflammation. **A.** Cecal LCN2 measurements at 60d after colonization in *Il10^-/-^* mice with full or reduced complexity synthetic microbiota as indicated in the “Colonized” line below each graph: Full SM14, just the SM10 species that do not degrade mucin *O*-glycans, or SM10 plus individual mucin degraders, *B. thetaiotaomicron* (*Bt*), *B. caccae* (*Bc*), *B. intestinhominis* (*Bi*) or *A. muciniphila* (*Am*) (n=7–23). **B.** Cecal histology scores of the same treatment group shown in A (n=5–20). **C.** Mucus thickness in SM10 colonized mice fed the FF diet (gray) compared to SM14 colonized mice fed either diet (n=5). **D.** Survival curves of *Il10^-/-^* adult mice colonized with SM14, SM10 and SM10+*Bt* to 150d. **E.** Regional histology of gastrointestinal tissues from SM10+*Bt*. **F.** Cecal LCN2 from SM14 and SM10+*Bt* colonized mice. **G.** Representative histology from SM10, SM10+*Bt* and SM10+*Am* at 60d. **H.** Survival curves of *Il10^-/-^* and DKO adult mice colonized with the SM14. **I.-L.** Individual endpoint weight loss (I.), Cecal LCN2 (J.), colon histology (K.) and cecal histology (L.) for SM14-colonized *Il10^-/-^Muc2*^-/-^ double knockout (DKO) mice fed the FR or FF diets (n=5-23). **M.** Colon histology in *Il10^-/-^*(right) and DKO (left) mice showing worse disease in DKO mice. **N.** Survival curves of mono-colonized DKO adult mice. **O-Q.** Cecal LCN2 (O.), colon histology (P.) and cecum histology (Q.) of mice shown in N. In panels A, B, I-K, O bold horizontal bar represents the mean and lighter error bars the S.E.M. In panels A-C, I-K, O two-way ANOVA and post hoc test with Original FDR method of Benjamini and Hochberg was used for statistics.

Adding back single mucin-degrading bacteria to the SM10 did not elicit the same level of LCN2 observed with the full SM14 at 60d, indicating that multiple species may act synergistically (**Figure 2A**). Despite a lack of weight loss (not shown) and lower LCN2 at 60d, the presence of either *B. thetaiotaomicron* or *A. muciniphila* as the sole mucin-degrader did elicit histological inflammation in the cecum at 60d that was statistically identical to the full SM14 (**Figure 2B**), indicating that weight loss and LCN2 phenotypes can in some cases be separate from histological damage. A longer, 150d experiment in which mucin-degrading *B. thetaiotaomicron* was added back to the SM10 revealed that it significantly accelerated lethal weight loss (**Figure 2D**) and caused histological inflammation that was worse in the cecum (**Figure 2E**). Thus, this commensal acts like a conditional pathogen in the compromised *Il10^-/-^*host. Interestingly, the elevated cecal histology in SM+*Bt* colonized mice also occurred without elevated LCN2 (**Figure 2F**). Finally, consistent with the idea that low dietary fiber promotes the general proliferation of mucin-degrading bacteria, mice colonized with SM10 plus a single mucin degrader all exhibited expansion of that mucin-degrading bacterium (**Figure S2J–O**). This contrasts with SM14-colonized mice, which only show expansion of *Akkermansia muciniphila* and *B. caccae* in response to low fiber (**Figure S2J**), implying that some mucin degraders out compete others.

The FR diet could suppress inflammation through mechanisms unlinked to its role in blocking mucin-degrading bacteria. As a separate test of the protective role for mucus in the context of the FR diet, we bred *Il10^-/-^Muc2^-/-^*double knockout (DKO) germfree mice, colonized them with the SM14 and fed either the FR or FF diets. These mice lost weight quickly and needed to be sacrificed regardless of diet (**Figure 2H**) and both groups exhibited severe inflammation (**Figure 2I-L**). Interestingly, with uniform elimination of MUC2, inflammation in DKO mice developed throughout the lower intestine but was more severe in the colon than in the cecum (**Figure 2K-M**). Since mucus is known to be thinner and patchier in the cecum compared to colon^24^, we interpret this to mean that inflammation occurs first in the cecum of SM14-colonized *Il10^-/-^* mice (*i.e*., with MUC2) because this is a site with high bacterial density and patchier mucus causing it to be where the mucus barrier fails first (note that small intestine also has thin mucus, but far fewer bacteria).

We further leveraged the DKO mice to measure how individual bacteria from different phyla (*e.g*., with different types of ligands for pattern recognition receptors) elicit inflammation in the context of IL-10 deficiency and deficient mucus barrier. In support of the conclusion that a variety of commensal bacteria can stimulate inflammation in this system, we observed that monocolonization with two different Bacteroidetes (*B. thetaiotaomicron* and *B. uniformis*), *A. muciniphila* (Verrucomicrobia) and *Eubacterium rectale* (Firmicutes) all resulted in strong inflammation, albeit once more with variations in LCN2 (**Figure 2N-Q**).

### Some human gut symbionts possess transferrable, pathogenic qualities between microbiomes

Because SPF mice fed the FF diet did not develop inflammation as severe as SM14-colonized mice (**Figure 1H,K**), we performed co-housing experiments in which pups born to SPF- and SM14-colonized mothers were mixed at weaning and exposed to each other’s microbiomes. Co-housing SM14-colonized *Il10^-/-^* pups with SPF mice prevented the weight loss phenotype observed in response to FF-feeding (**Figure 3A**). However, it did not reduce—and even slightly increased—cecal inflammation as measured by LCN2 (**Figure 3B**) or histology (**Figure 3C**), again indicating that weight loss and inflammation can be uncoupled in the presence of different microbiota. Co-housed mice born to SPF mothers showed a more variable response , with ∼50% of these mice developing inflammation that was similar in severity to FF-fed, SM14-colonized mice (**Figure 3B,C**). Time course analysis of fecal LCN2 revealed a larger difference between co-housed and non-co-housed SPF mice, with co-housed SPF mice behaving nearly identically to FF-fed SM14 (**Figure 3D**, solid purple).

**Figure 3.**
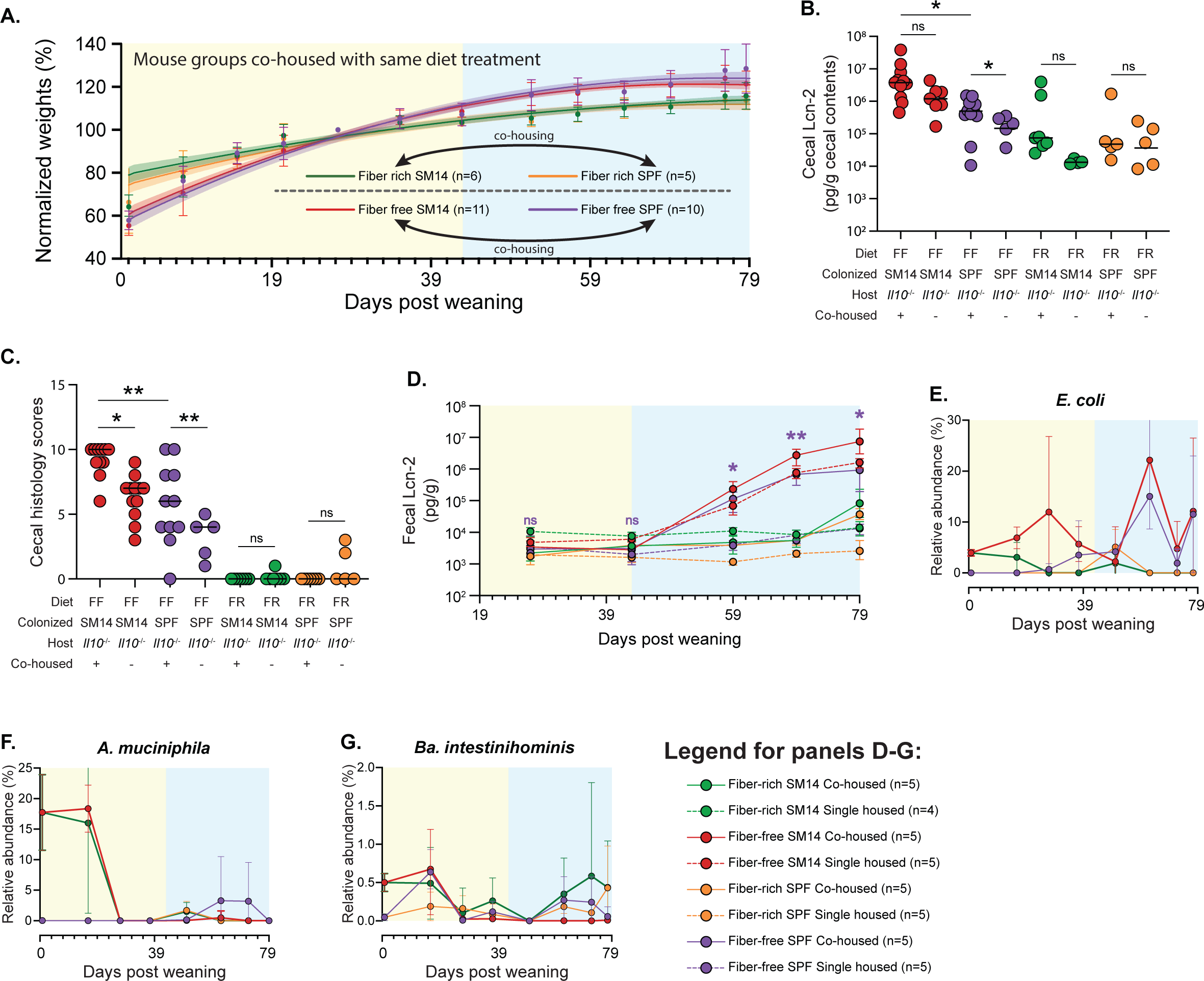
Co-housing SPF- and SM14-colonized mice after weaning worsens disease. **A.** Weights of SPF or SM14-colonized mice co-housed at weaning (21d) with pups harboring the opposite colonization. **B.** Cecal LCN2 levels in co-housed or non-co-housed SPF and SM14 mice (co-housing status is indicated at the bottom of this and other panels; n=5-11, one-tailed Student’s t-test and Wilcoxon test). **C.** Cecal histology of the same treatments shown in B (n=5-11, one-tailed Student’s t-test and Wilcoxon test). **D.** Cecal LCN2 over time in co-housed and non-co-housed SPF and SM14 mice. **E–G.** Relative abundances of select SM14 species invading SPF co-housed mice, *E. coli*, *A. muciniphila*, *Ba. intestinihominis*, respectively. Data are represented as mean ± SEM. *p < 0.05; **p < 0.01; ***p < 0.001; ****p < 0.0001.

Based on 16S rRNA gene sequencing, at least 6 members of the SM14 could be detected in co-housed SPF mice, often transiently or near the end of the time course when inflammation developed (**Figures 3E-G, S3A-I**). The most prominent of these invading bacteria was the human commensal *E. coli* strain HS, which appeared in co-housed SPF mice around 28 dpw, prior to the onset of inflammation, and gradually increased, eventually reaching levels >10% in most mice (**Figure 3E**, solid purple). Two of the mucin-degrading SM14 bacteria (*A. muciniphila* and *Ba. intestinihominis*) showed a similar trend, albeit reaching lower levels and with variability among individual mice (**Figure 3F,G**). Two other bacteria (*C. aerofaciens* and *B. uniformis*) showed transient increases around the time inflammation was increasing (∼48 dpw) and these organisms decreased thereafter (**Figure S3D,E**). A test of whether weekly gavages of *E. coli* HS into SPF mice that were otherwise not exposed to SM14 bacteria failed to support the hypothesis that this species is the sole cause of increased inflammation (**Figure S3J,K)**. Taken together, these co-housing results demonstrate that some of the 14 human bacteria in our synthetic microbiota are capable of transferring increased low fiber-induced inflammation to mice colonized with a natively complex microbiota, extending the relevance of this diet-driven model to more complex communities and implying that one or more of the SM14 bacteria is endowed with conditional pathogenic qualities.

### Certain gut bacteria and metabolites associated with exclusive enteral nutrition prevent inflammation despite mucus erosion

A notable characteristic of the disease-promoting FF diet is that its macronutrient composition resembles some exclusive enteral nutrition (EEN) diets used clinically to treat some IBDs (**Table S1**). EEN diets often contain low or no fiber^25^ and have proven to be effective at inducing remission in humans, although precise mechanisms are still unknown^26^. To determine if an EEN diet lacking fiber promotes inflammation in our gnotobiotic *Il10^-/-^* model, we weaned SM14-colonized pups onto a commercial EEN diet, which is normally taken as a liquid but in this case was freeze-dried, pelleted and sterilized. The average weight trajectories of 15 mice supported the conclusion that the low fiber EEN diet promotes some weight loss, although not as severe as the FF diet (**Figure 4A**). Cecal LCN2 measurements and histology revealed substantial disease variation in individual animals, with some mice resembling healthy FR-fed mice, some resembling diseased FF-fed mice and some intermediate (**Figure 4B,C**). Despite experiencing less inflammation, the EEN mice still exhibited reduced mucus thickness, which we expected given the lack of fiber in the formula used (**Figure 4D**). Measurements of short- and branched-chain fatty acids (SCFAs, BCFAs) revealed that mice fed the EEN diet displayed between 43.2-723.0-fold (mean 133.4) elevated amounts of a single BCFA, isobutyrate (**Figure 4E**). Suggesting that higher production, might help suppress inflammation.

**Figure 4.**
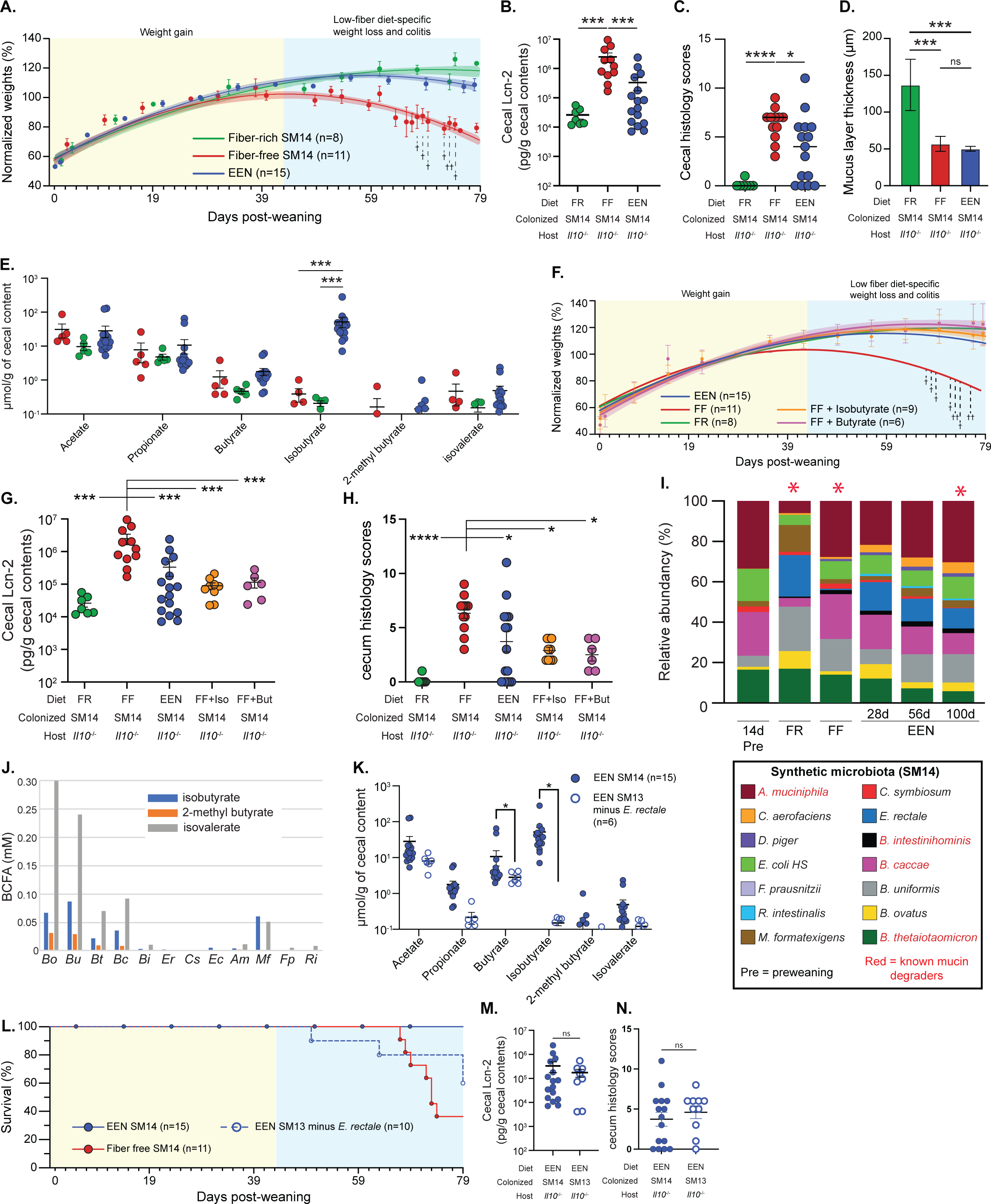
Fiber free EEN diet improves inflammation in part through isobutyrate production. **A.** Weights of mice weaned onto FR, FF, and EEN diets. **B.** Cecal LCN2 levels in mice weaned onto FF, FR, and EEN diets (n=7-15). **C.** Cecal histology scores of the mice shown in B. (n=7-15). **D.** Mucus thickness measurements in mice shown in B.-C. (n=5). **E.** Short- and branched-chain fatty acid measurements in mice fed FF, FR, or EEN diets (n=5-15). **F.** Weights of mice fed the FF diet and water supplemented with either 35mM isobutyrate (orange) or butyrate (purple) (n=6-15). **G.** Cecal LCN2 levels in the mice from panel F. **H.** Cecal histology scores of the mice shown in F. (n=6-15). **I.** SM14 community composition in mice fed the FR, FF and EEN diets. Red asterisks denote FF and EEN diet comparison. **J.** Short- and branched-chain fatty acid measurements in culture supernatants of indvidual SM14 bacteria. **K.** Short and branched chain fatty acid measurements in EEN fed mice colonized with either the full SM14 or an SM13 lacking *E. rectale* (n=6-15, one-tailed Student’s t-test and Wilcoxon test). **L.** Survival of SM14 and SM13 (minus *E. rectale*) colonized mice on the FF and EEN diets. **M.** Cecal LCN2 measurements of the mice shown in L. (n=6-15, one-tailed Student’s t-test and Wilcoxon test). **N.** Cecal histology scores of the mice shown in L. (n=10,15). Data are represented as mean ± SEM. *p < 0.05; **p < 0.01; ***p < 0.001; ****p < 0.0001. In panels B, C, D, E, G, H, two-way ANOVA and post hoc test with Original FDR method of Benjamini and Hochberg was used for statistics.

Isobutyrate is an isomer of the well-studied, anti-inflammatory short-chain fatty acid butyrate, which did not increase (**Figure 4E**), and isobutyrate is derived from L-valine fermentation by certain gut bacteria^27^. Two other BCFAs (2-methyl butyrate and isovalerate) did not increase despite the amino acids they are derived from (L-leucine and L-isoleucine) being present at similar abundance^28^ in the soy and milk protein used in the EEN formulation (**Figure 4E**). This suggests that the increase in isobutyrate is not attributable to increased bulk dietary protein fermentation, which should increase all three BCFAs. Providing either isobutyrate or butyrate (35 mM) in the drinking water of mice fed the disease-promoting FF diet decreased weight loss (**Figure 4F**) as well as inflammation (**Figure 4G,H**), revealing that either of these molecules can offset the diet- and microbiome-induced damage.

EEN feeding promoted significant changes in the SM14, most notably 211-278-fold increases (mean 255.6) in *E. rectale* relative abundance (**Figure 4I**, compare columns with red asterisks). While this Firmicute is known to produce butyrate, measurements of culture supernatants from 13 of the SM species in medium supplemented with L-valine revealed that *E. rectale* does *not* produce isobutyurate under the conditions tested. Rather, isobutyrate was produced by all 4 of the *Bacteroides,* plus *M. formatexigens* (**Figure 4J**). Nevertheless, to determine if the large increase in *E. rectale* abundance was functionally connected with isobutyrate production, we bred neonatal mice born to parents harboring a version of the SM14 but lacking *E. rectale* (“SM13 minus *E. rectale*”) and fed them the EEN diet. Consistent with a causal role for *E. rectale* in isobutyrate production, mice lacking *E. rectale* failed to produce isobutyrate (**Figure 4K**). These mice also exhibited 40% lethality at 79dpw (**Figure 4L**), although there was not a significant increase in cecal LCN2 levels or histology relative to the EEN/SM14 group (**Figure 4M,N**).

### Mucin-degrading bacteria mediate fiber deprivation-induced immune responses in a time- and location-dependent manner

Development of intestinal inflammation is a complex process involving innate and adaptive immune responses and critical roles for Th1/Th17 cells have been described in the *Il10^-/-^* colitis model in conventional/SPF mice^19^. Nevertheless, how specific microbial triggers influence the underlying immune pathways and how responses develop over time are less clear. We repeated the same SM14 and SM10 (non-mucin-degrading community) colonization experiments described above with a different C57BL/6J line of *Il10^-/-^* mice (mouse facility: University of Luxembourg) and observed similar weight loss in both adult and post-weaning FF-fed mice colonized with SM14, but not SM10 (**Figure 5A-C**). Given the temporal variation in colitis phenotypes in FF-fed SM14- (faster colitis) and SM10-colonized (slower colitis) *Il10^-/-^* mice, our model provides opportunities to investigate the contributions of not only the microbial triggers (*i.e*., with or without mucin degraders) but the regionality (cecum vs. colon) of inflammatory pathways associated with increased microbial mucin foraging during fiber deficiency.

**Figure 5.**
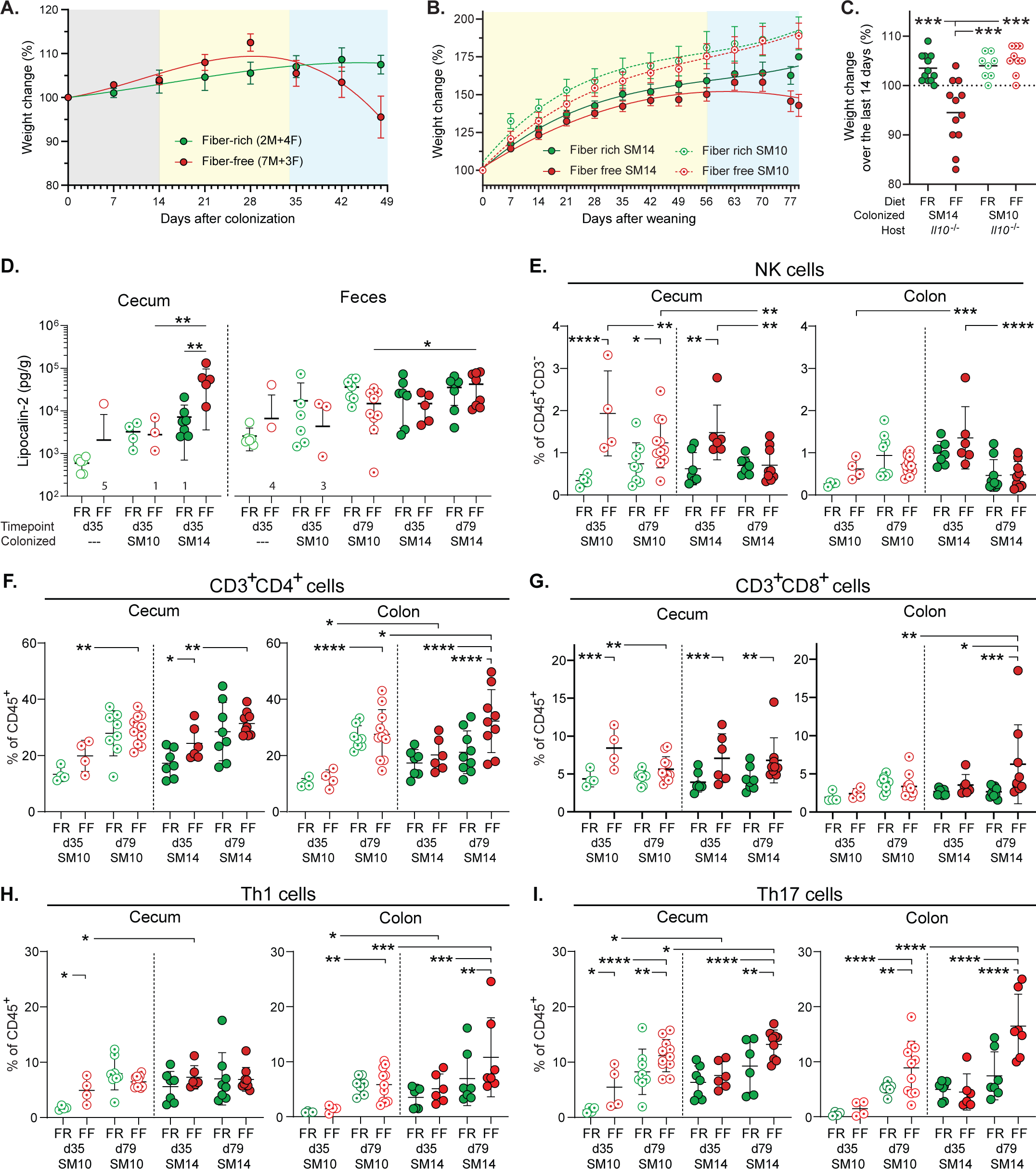
Mucin-degrading bacteria mediate low fiber-induced immune responses in a time- and location-dependent manner. **A.–B.** Weights of SM14-colonized adult *Il10^-/-^* mice (**A.,** n=6–10) and *Il10^-/-^* pups born to SM14- or SM10-colonized dams (**B.,** n=8–12), reproduced in the University of Luxembourg mouse facility. **C.** Endpoint weights of individual animals shown in B. (n=8–12). **D.** LCN2 levels in the cecal contents and feces of GF and colonized *Il10^-/-^* mice (n=4–7). Numbers on the X axis indicate the number of samples in which LCN2 was not detected. **E.** Proportion of NK cells among CD3^-^CD45^+^ cells in the cecum and colon lamina propria of SM10- and SM14-colonized *Il10^-/-^* mice (n=4–11). **F–I.** Proportion of CD3^+^CD4^+^ (**F.**), CD3^+^CD8^+^ (**G.**), Th1 (**H.**) and Th17 (**I.**) cells among CD45^+^ cells in the cecum and colon of SM10- and SM14-colonized *Il10^-/-^* mice (n=4–11). Helper T (Th) cell subsets are defined as CD3^+^ CD4^+^ Foxp3^-^. Data are represented as mean ± SD. *p < 0.05; **p < 0.01; ***p < 0.001; ****p < 0.0001. In panels C-I, two-way ANOVA and post hoc test with Original FDR method of Benjamini and Hochberg was used for statistics.

In FF-fed SM14-colonized mice, increased LCN2 was detected in the cecum as early as 35 dpw compared to FR and this diet effect was not observed in mice colonized with SM10 or left germfree (**Figure 5D**). SM14 mice weaned onto the FF diet also exhibited higher fecal LCN2 (a proxy for colon) than SM10 mice at 79 dpw, supporting a pro-inflammatory role of mucin-degrading bacteria during fiber deprivation (**Figure 5D**). An expansion of natural killer (NK) cells was detected in the cecum of FF-fed mice as soon as 35 dpw and this expansion was similar in SM10- and SM14-colonized mice (**Figure 5E**, left panel; see **Figure S4A** for flow cytometry sorting scheme and **S4B-I** for germfree *Il10*^-/-^ and SM14-colonized WT controls for flow cytometry experiments). In contrast to the pattern observed in cecum, NK cells expanded in the colons of both FR and FF-fed SM14-colonized mice as soon as 35 dpw, but not in the colons of SM10-colonized mice (**Figure 5E**, right panel). This further suggests that mucin-degrading bacteria are more important to elicit responses in the colon where mucus is thicker and degradation is required to increase contact. We also observed region-specific host responses that are driven by diet x host genotype interactions, since FF-fed GF *Il10^-/-^* mice, but not SM14 or SM10 colonized WT mice, showed increased NK cells in the cecum, but not in the colon, at 79 dpw compared to FR-fed controls (**Figure S4B**).

While NK cells were more abundant at 35 days and later decreased, T cell recruitment increased over time, generally reaching highest levels at 79 dpw, in both the cecum and colon of SM10- and SM14-colonized *Il10^-/-^*mice (**Figures S5A, S4C**). In the ceca of both SM10- and SM14-colonized *Il10^-/-^* mice, the FF diet increased CD4^+^ and CD8^+^ T cell populations by 35 dpw, with CD4^+^ T cells increasing further by 79 dpw independently of the diet (**Figure 5F,G**). However, CD4^+^ T cells also accumulated over time in the ceca of FF-fed GF *Il10*^-/-^ mice (**Figure S4D**), while CD8^+^ T cells were induced by FF-feeding in the ceca of GF *Il10*^-/-^ mice, as well as in the ceca and colons of SM14-colonized WT mice (**Figures S4E**). These results suggest that fiber deprivation induces the recruitment of T cell populations through multiple paths and not all of these are regulated by mucin-degrading bacteria, the microbiota and IL-10 deficiency.

Consistent with previous descriptions of *Il10^-/-^* colitis in conventional mice^19^ both SM10- and SM14-colonized mice generally exhibited infiltration of Th1 and Th17 cells over time (**Figure 5H,I**) with fewer changes in Th2 cells (**Figure S5B**). Especially for Th17 cells, this infiltration was higher in FF-fed mice in both the cecum and the colon (**Figure 5I**). While Th17 cells reached nearly equivalent levels at 79 dpw in SM10 and SM14-colonized ceca, overall levels were lower in the SM10-colonized colons compared to SM14 (**Figure 5I**). Furthermore, the FF diet increased Th1 levels in the SM14-colonized colons, but not in the SM10-colonized ones (**Figure 5H**). Consistent with the readouts mentioned above, this observation supports the idea that the role of mucin-degrading bacteria is more important in the colon where erosion of mucus facilitates the induction of the anti-microbial Th1 responses.

Cytokine measurements in cecal tissues revealed expected increases in IBD-associated markers IL-1β, IL-6, IL-17, IL-22, TNF-α and IFN-γ in fiber-deprived *Il10^-/-^* mice that were colonized with SM14 as adults (**Figure 6A-F**).To provide functional support to the immune cell profiles, we compared levels of Th1- and Th17-type cytokine transcripts in ceca and mesenteric lymph nodes at 35 and 79 dpw in mice colonized at birth. In SM14-colonized *Il10^-/-^*mice, the FF diet led to increased expression of Th1 (IFN-γ, IL-6, TNF-α, IL-12) and Th17 cytokines (IL-17F, IL-22, IL-23 and TGF-β), as well as the mucin-inducing cytokine IL-13 (**Figures 6G-U**). Interestingly, in the ceca of SM14-colonized mice, both Th1 and Th17 cytokines were induced at 35 dpw, while only Th1 cytokines were maintained at 79 dpw (**Figure 6G-N**). These cytokine responses tended to develop more slowly in SM10-colonized ceca compared to those with SM14 and, both Th1 and Th17 cytokines were increased by 79 dpw in FF-fed SM10-colonized mice (**Figure 6G-N**). In the colon-draining mesenteric lymph nodes (MLNs) where naïve T cells are activated and polarized, the Th17-related cytokines, IL-17F and IL-22, were increased by FF-feeding compared to FR in both SM10- and SM14-colonized mice (**Figure 6S-T**), while the Th1-polarizing cytokines IFN-γ, IL-6 and IL-12 were only increased in SM14-colonized mice, consistent with a dependence on mucin-degrading bacteria to develop Th1 responses (**Figures 6P-R**).

**Figure 6.**
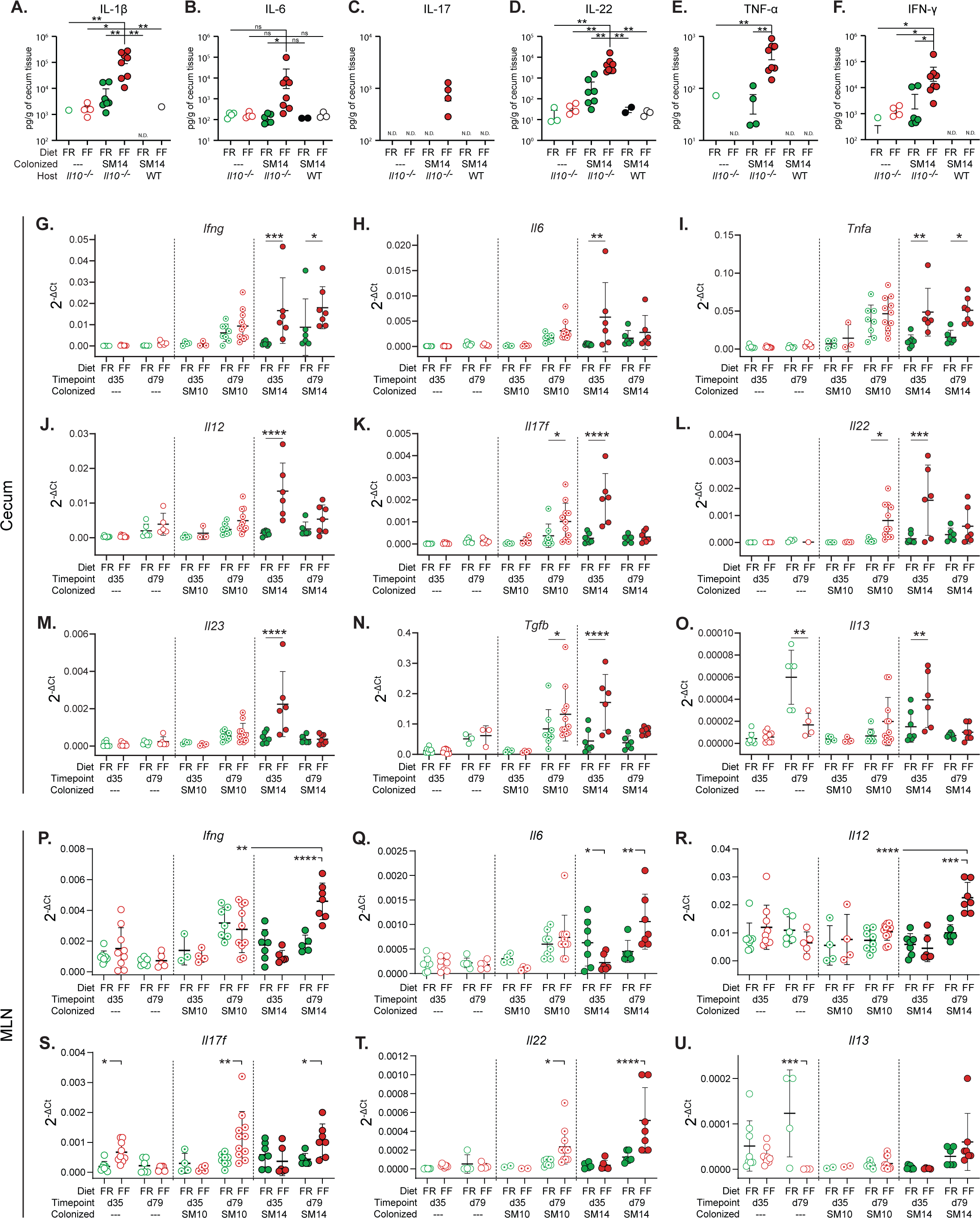
Mucin-degrading bacteria mediate fiber deprivation-induced Th1 inflammation. **A.–F.** Cytokine measurements at 60d (or earlier in SM14/FF mice that lost >20% weight) by Luminex bead assay in the cecum of wild-type and *Il10*^-/-^ mice colonized as adults. **G.–U.** Cytokine mRNA expression in the cecum (**G.–O.**) and Mesenteric Lymph Nodes (**P.–U.** MLN) of *Il10*^-/-^ pups weaned to FF or FR diets (n=4-11, two-way ANOVA and post hoc test with Original FDR method of Benjamini and Hochberg). Data are represented as mean ± SD. *p < 0.05; **p < 0.01; ***p < 0.001; ****p < 0.0001.

Despite the IL-10 deficiency, Foxp3^+^ regulatory T cell (Treg) recruitment was also higher during FF feeding of SM14 *Il10^-/-^*mice (**Figures S5C-F**). This expansion being independent of IL-10 suggests a mechanism driven by other regulatory mediators such as TGF-β, whose transcript levels increased in SM14-colonized cecal tissues as soon as 35 dpw (**Figure 6N**). Regulatory T cells expressing Tbet or RORγt have been proposed as counter-regulators of inflammatory Th1 and Th17 cells, respectively^29,30^. Consistent with this, the abundance of Tbet^+^ Treg cells follow the same trends as inflammatory Th1 cells with higher levels in FF-fed, SM14-colonized mice (**Figures S5D**). By contrast, the highly suppressive RORγt^+^ Treg population was reduced at 35 dpw in the colon and at 79 dpw in the cecum of FF-fed, SM14-colonized *Il10^-/-^* mice (**Figure S5E**), thus favoring inflammatory responses. In addition, Gata3^+^ Tregs were increased in FF-fed colonized colons (**Figure S5F**). While their role in the specific regulation of type 2 inflammatory cells is still unclear, they may constitute a reservoir of Tregs required for tissue repair by 79 dpw in colonized colons^31,32^. Finally, despite the general expansion of Treg, early (35 dpw) loss of the highly suppressive RORγt^+^ subset and the IL-10 deficiency are likely to allow the Th1/Th17 responses to flourish in FF-fed, SM14 mice. Together, these results reveal strong Th1/Th17 immune pathways induced by fiber deprivation, differentially regulated by microbial community members and host genetics, in a time- and intestinal site-specific manner.

### Alterations to IgA–microbiota interactions precede inflammation

IgA production is a common anti-microbial response in the gut and is usually upregulated during colitis in humans and mouse models^33^. Mirroring the LCN2 trend observed for early inflammation (**Figure 6A**), soluble IgA titers were increased in the cecum at 35 dpw and the feces at 79 dpw, in FF-fed SM14-colonized *Il10^-/-^* mice compared to FR, but not in SM10-colonized mice (**Figure 7A**). However, prolonged FF-feeding (79 dpw) resulted in depletion of plasma and IgA-producing cells in both the cecum and colon and this loss was also observed in FF-fed WT mice that were colonized with SM14 (**Figure S6A, B**), suggesting a diet-driven effect on IgA-producing cells rather than an effect of IL-10 loss. This is surprising considering the increased levels of secreted IgA (**Figure 7A**), but consistent with a published report showing reduced titers of serum IgA and fewer IgA^+^ B cells in the small intestine of wild-type mice fed a zero-fiber diet compared to high-fiber^34^. In parallel with reduced IgA-producing cells after FF-feeding, the proportion of IgG-producing B cells was increased at 35 and 79 dpw in both the cecum and colon of SM14-colonized *Il10^-/-^* mice (**Figure S6C,D**). Consistent with the high production of Th1 cytokines in the presence of mucin-degrading bacteria, FF-feeding increased the proportion of IgG-producing cells only in SM14-but not in SM10-colonized colons (**Figure S6D**)^35^.

**Figure 7.**
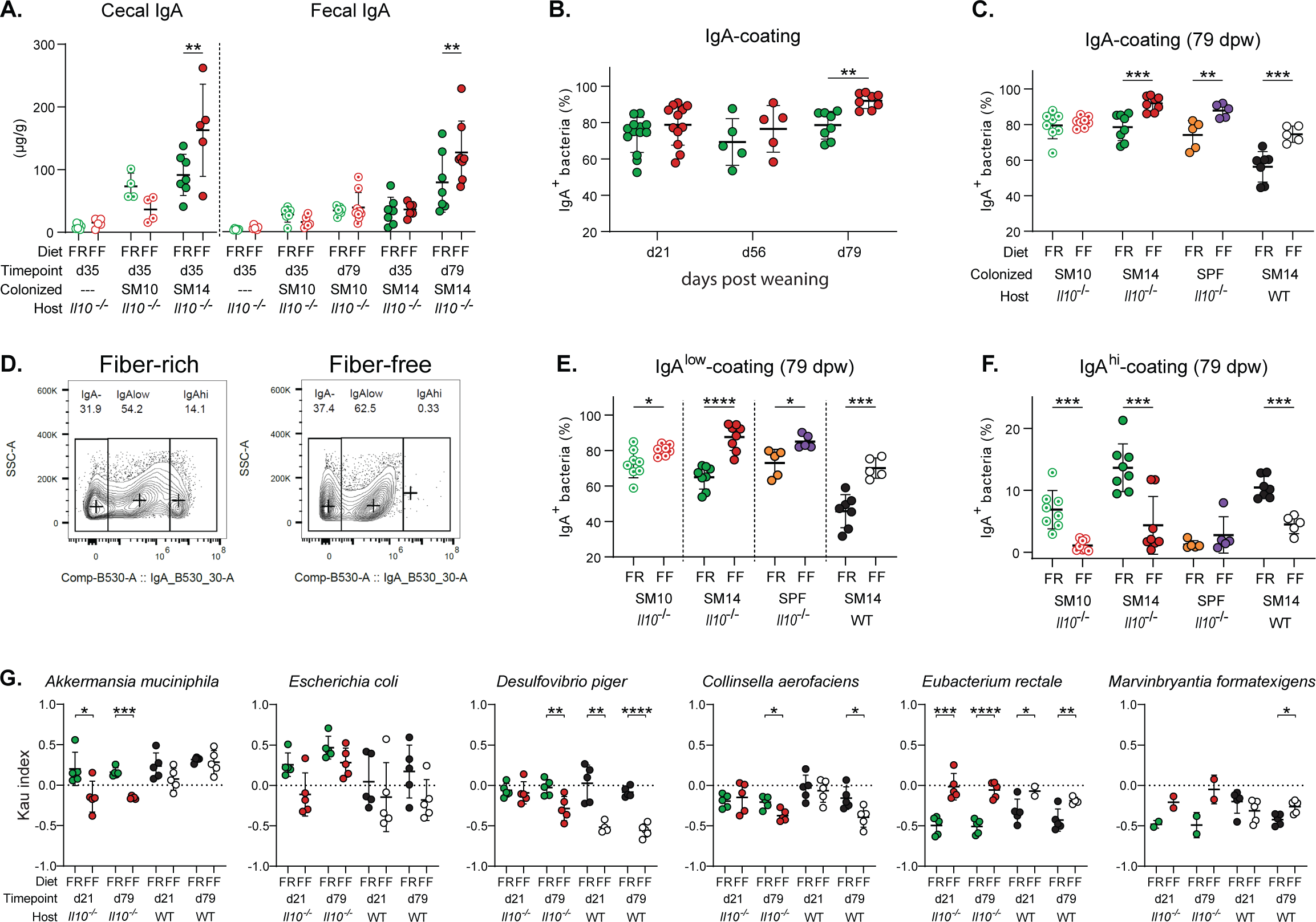
Fiber deprivation alters IgA–bacteria interactions. **A.** Concentrations of free IgA in the cecal contents and feces of GF and colonized *Il10^-/-^* mice (n=4–9). **B.** Percentages of total IgA-coated bacteria in the feces of FR- and FF-fed, SM14-colonized mice at 21, 56 and 79 dpw (n=5–13). **C.** Percentages of total IgA-coated bacteria in the feces of SM10-, SM14- or SPF-colonized *Il10^-/-^* mice and SM14-colonized WT mice fed the FR or the FF diet for 79 days (n=5– 8). **D.** IgA-coating profiles of fecal bacteria from SM14-colonized mice fed a FR (left) or a FF (right) diet for 56 days showing the gating strategy of populations being low-coated (IgA^low^) and high-coated (IgA^high^). Total IgA coating consists of the addition of IgA^high^ and IgA^low^ coating. **E–F.** Percentages of IgA^low^- (**E.**) and IgA^high^- (**F.**) coated bacteria in the feces of indicated groups at 79 dpw (n=5**–**8). **G.** IgA-coating index (Kau index) of fecal bacteria from SM14-colonized *Il10^-/-^* and WT mice (n=2–5, multiple unpaired t-test and post hoc test with the two-stage linear step-up procedure of Benjamini, Krieger and Yekutieli). Data are represented as mean ± SD. *p < 0.05; **p < 0.01; ***p < 0.001; ****p < 0.0001. In panels A., B., C., E., F., two-way ANOVA and post hoc test with the two-stage linear step-up procedure of Benjamini, Krieger and Yekutieli was used for statistics.

Since coating with immunoglobulin A (IgA) has been proposed to identify bacteria that are potentially more colitogenic^36^, we next focused on how the inflammation-promoting FF diet alters bacterial IgA coating in *Il10^-/^*^-^ mice. Consistent with previous studies^33^, the amount of total IgA-coated bacteria was higher in FF-fed mice than in FR-fed mice at 79 dpw (**Figure 7B**). Along with increased luminal IgA (**Figure 7A**), this FF diet-induced increase in total IgA-coating was not observed in SM10-colonized mice but was observed in SPF *Il10^-/-^* mice and SM14-colonized WT mice, supporting a mechanism driven by mucin-degrading bacteria (**Figures 7C, S7A**). Intriguingly, analysis of IgA-coated bacteria revealed the presence of 2 differentially coated populations in FR-fed mice: a large population with low-coating and a smaller population with high coating, and both populations responded differently to FF-feeding (**Figure 7D**). The proportion of low-coated bacteria increased in FF-fed mice, while the highly-coated bacteria were almost completely lost in FF-fed mice, as early as 21 days of feeding (**Figure 7D-F, S7B**). Additionally, highly-coated bacteria were diminished in SM10-colonized mice as in SM14-colonized mice as soon as 21 dpw and low-coated bacteria only slightly increased by 79 dpw (**Figure 7E,F, S7C**). Together these results suggest two levels of regulation of IgA coating by the FF diet: a rapid loss of the high coating that occurs in the absence of mucin-degrading bacteria, and a later increase of low and total coating that is partially dependent on the mucin-degrading bacteria and likely linked to the increased secretion of IgA. Interestingly, the EEN diet that dampened colitis but not mucus thinning in SM14-colonized *Il10^-/-^* mice, increased the proportion of IgA-coated bacteria as soon as 21dpw and conserved the high-coated population compared to mice fed FF (**Figure S7D**), suggesting additional immune-regulatory properties of EEN diet components.

Given the altered pattern of overall IgA coating, we examined IgA coating of individual SM14 bacteria. Consistent with the early loss of highly coated bacteria, changes in IgA-coating of SM14 members also appeared as soon as 21 days of FF-feeding (**Figures 7G, S7E**). Among the SM14 members, *A. muciniphila*, *D. piger*, *E. coli* and *C. aerofaciens* showed reduced IgA coating index (ICI) values at one or both timepoints in FF-fed *Il10^-/-^*mice (**Figures 7G**). In WT mice, this FF diet-induced reduction in IgA coating was less severe for *A. muciniphila*, but more pronounced for *D. piger* and *C. aerofaciens*, revealing that IL-10 deficiency affects IgA-coating of commensals in a species-dependent manner. In contrast, *E. rectale* showed a low ICI in FR-fed mice and this increased with FF-feeding (**Figure 7G**), a condition in which it is present at very low levels. The causes and effects of variations in IgA coating are still being determined. However, our results demonstrate that different combinations of microbial colonization, diet and host immune status can alter both the amount of intestinal IgA and the bacteria it targets. Along with the data shown above, our results suggest the possibility that fiber deprivation initiates early disruptions in IgA–microbiota interactions along with a loss of IgA-producing plasma cells. Additional experiments to sort out the respective contributions of altered IgA coating vis-à-vis alterations in microbiota physiology are made possible by these observations in a tractable gnotobiotic system.

## Discussion

The pathophysiology of IBDs is complex and variable, in part because of the large number of different genetic contributions that combine with environmental, microbial and dietary triggers known or hypothesized to influence its development. The diet-driven inflammation model investigated here provides both a case study, in which contributing factors to *Il10^-/-^*-associated inflammation can be explored at mechanistic levels, as well as a more general experimental paradigm in which host genetic, microbiota and dietary variables can be manipulated to examine their effects on disease development. As examples of spontaneous, genetically driven intestinal inflammation continue to be identified^37–39^, the ability to work with these murine models in germfree/gnotobiotic conditions and with defined diets holds the potential to uncover foundational principles about these complex diseases.

An important conceptual idea that is supported by our findings is that opposing bacterial functions influence disease outcomes. Low fiber induced mucus erosion is deleterious, but mostly in the context of some members of the human SM14 community that appear to have pathogenic qualities that can transfer their pro-inflammatory effect to SPF mice. During EEN feeding, mucus erosion still occurs, but is offset to some degree by production of isobutyrate. These opposing functions do not need to be mechanistically connected; rather, they may work independently as long as the protective function (isobutyrate) is capable of reducing inflammation that results from mucus thinning. Isobutyrate production is both influenced by diet and contingent upon *E. rectale*. Thus, these data present actionable hypotheses for future work in humans to determine if patients that respond positively to EEN also show increased isobutyrate and if EEN responses can be stratified based on presence of *E. rectale*. Deconstruction of the EEN formula to identify ingredients that promote *E. rectale*-dependent isobutyrate production, as well as identifying the bacteria that produce isobutyrate *in vivo*, may also lead to optimization of EEN formulations. These studies are ideally suited to be first conducted in a simplified pre-clinical model that allows focus on individual contributions prior to extending to more complex human studies.

The link between low dietary fiber and compromised thickness and/or permeability of the mucus layer has been emerging through several studies^13,14,17,15,16^. Here, we make direct functional connections between the dietary fiber-gut microbiome-mucus layer axis in the context of IBD development in a genetically susceptible host in a way that depends on the specific bacteria present (SM14 vs. SPF). Many questions remain regarding the development of the multifactorial diseases called IBDs. While host genetics are a mostly permanent trait that is difficult to repair, an exception being stem cell transplant in children lacking IL-10 signaling^40^, the microbiome and especially diet are factors that could be manipulated to delay or reverse disease. Ever since the discovery that rearing *Il10^-/-^*mice under germfree conditions abrogates inflammation^20^, there has been substantial interest in finding specific bacteria that elicit inflammation when added back to these animals. This search has yielded insight into the contributions of bacteria with known pathogenic qualities, such as *Helicobacter* spp., *Enterococcus faecalis* and adherent invasive *E. coli*^19^. However, roles for specific commensal bacteria have also been established using gnotobiotic mice^41^, albeit without a clear understanding of the mechanism(s) involved. Based on data presented here, we propose that it is appropriate to focus more on the influence of positive and negative bacterial functions than on the role of specific taxa. The rationale for this is that metabolic pathways like mucin degradation can be present in diverse bacteria and some of these organisms may fulfill similar roles in eroding mucus, while some may not. In contrast, strains of the same species can vary in key metabolic capabilities, including mucus degradation, which is known to vary substantially among strains of *Bacteroides*^42^. If one considers the effects of the microbiome on IBD development to be a cumulative series of positive and negative stimuli caused by the particular behaviors and metabolites exhibited by the microbes present, it should be possible to optimize beneficial processes (*e.g*., butyrate, isobutyrate) while reducing detrimental events like mucus erosion. While some of these levers can be manipulated through diet, a promising path for intervention may eventually include adding or replacing specific bacteria within a person’s individual microbiota with more optimal strains or perhaps even those that are engineered to increase beneficial metabolites.

## Acknowledgements

We thank the University of Michigan Germfree and Microbiome Cores for technical assistance, Leonard Augenlicht for Muc2^-/-^ mice and Bill Krueger (J. Rettenmaier) for fiber samples. We are grateful for support from the Kenneth Rainin Foundation (Innovator Award, ECM) and the US NIH (R01s DK118024, DK125445 to ECM; P01 HL149633 funding to CAL, TMS and ECM). We thank the Luxembourg National Research Fund (FNR) for CORE grants (C15/BM/10318186, C18/BM/12585940) and BRIDGES grant (22/17426243) to MSD and MEDICE Arzneimittel Pütter GmbH & Co. KG, Germany and Theralution GmbH, Germany for funding through the public–private partnership FNR BRIDGES grant to MSD (22/17426243). MB was supported by a European Commission Horizon 2020 Marie Skłodowska-Curie Actions individual fellowship (897408). MW was supported by a Fulbright grant for Visiting Scholars from the Commission for Educational Exchange between the USA, Belgium and Luxembourg. ETG was supported by FNR PRIDE (17/11823097). CA was supported by the University of Michigan Postdoctoral Translational Scholars Program.

## Competing interest statement

ECM works as a consultant and an advisory board member at January, Inc, United States. MSD works as a consultant and an advisory board member at Theralution GmbH, Germany.

## Supplemental Figure Legends

**Supplemental Figure 1.**
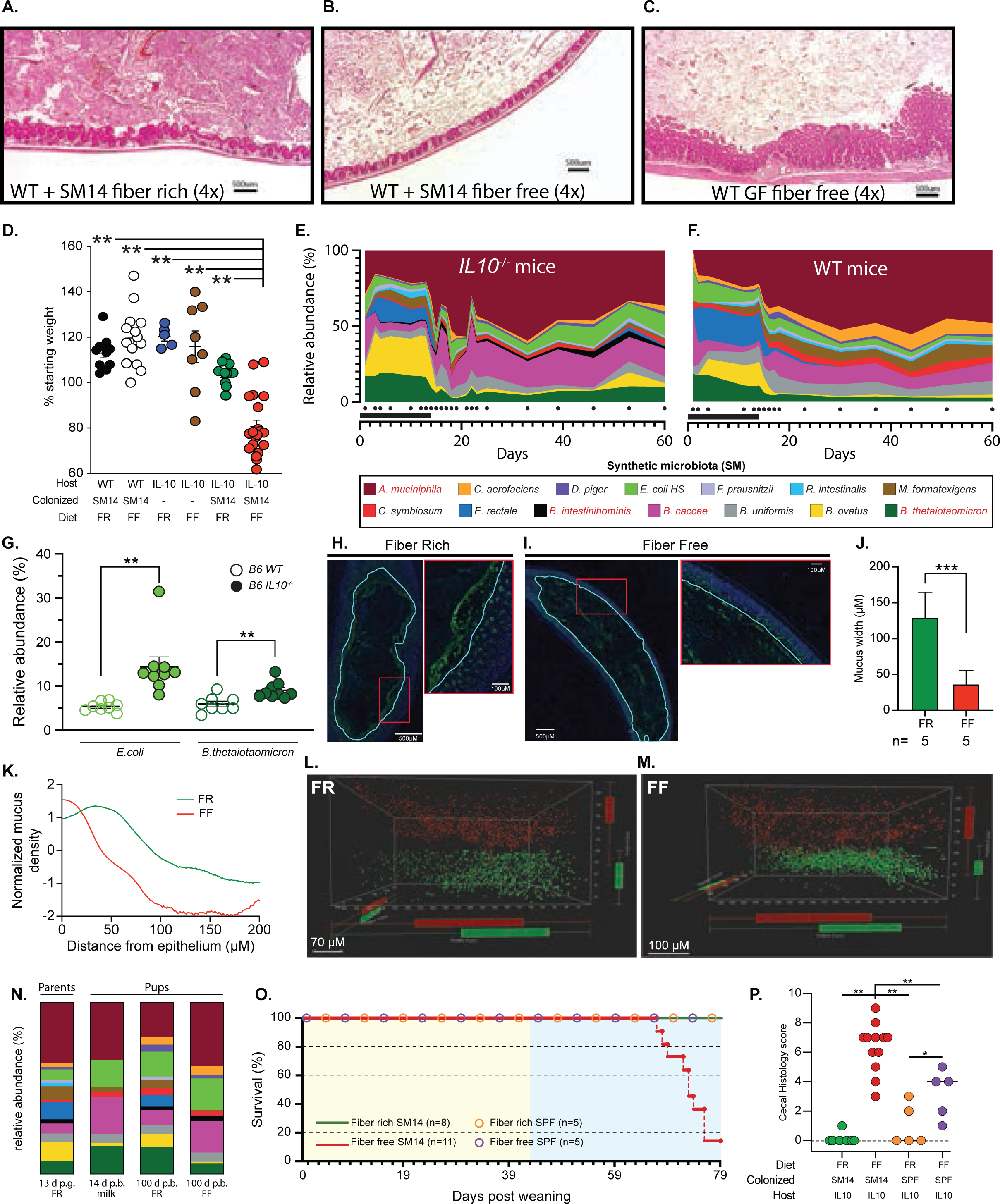
Low fiber and microbiome driven inflammation development in *Il10^-/-^* mice. **A.-C.** Representative histology from wild-type (WT) SM14-colonized mice fed FR (A.), WT SM14-colonized mice fed FF (B.) or WT germfree mice fed FF (C.). All pictures are shown at 4x magnification, bars 500μM. **D.** Endpoint weights of the mice shown in Figure 1A main text. Bold horizontal bar represents the mean and lighter error bars the S.E.M. (n=7–20) **E.–F.** Relative abundance plots of SM14 bacteria in *Il10^-/-^* (E.) or wild-type (F.) mice shifted to the FF diet at 14d after colonization. Dots below the graph indicate the time points at which feces was sampled and the black bar on the x-axis represents the period of FR feeding. **G.** Comparison of cecal *E. coli* and *B. thetaiotaomicron* populations in *Il10^-/-^* vs WT mice after 60d of colonization. Bold horizontal bar represents the mean and lighter error bars the S.E.M. (n=8-9, one-tailed Student’s t-test and Wilcoxon test). **H.–I.** Representative staining and imaging for Muc2 (green) in colonic sections from FR (H.) or FF (I.) and counter-stained with DAPI (blue). These images were used for automated mucus measurement using BacSpace software^14^ as described in methods. **J.** Automated mucus thickness measurements generated from colonic section images like those shown in H.**–**I. (n=5, one-tailed Student’s t-test and Wilcoxon test). Five mice were measured per treatment, with a total of 1**–**2 fecal mass sections imaged per mouse. **K.** Averaged Muc2 staining intensity measurements for the mice represented in H.-J. as a function of distance from the epithelium. **L.–M.** Representative visualization of 1 um-sized beads (in red) layered on the distal colon epithelium (in green) after feeding FR (L.) or FF (M.) to SM14-colonized *Il10^-/-^*mice. The space between the beads and epithelial cells represents the penetrability of the mucus layer. **N.** Representative SM14 composition in adult mice fed FR at 13d post gavage (p.g.) compared to still suckling pups at 14d post birth (p.b.) and 100d old SM14 colonized pups that were weaned onto either FR or FF diets. **O.** Survival plot of SM14 or SPF colonized pups weaned onto FR or FF diets and maintained for a total of 100d, the typical experiment duration. **P.** Cecal histology scores for SM14 or SPF colonized pups weaned onto FR or FF diets and maintained for a total of 100d. P. Cecal histology scores for SM14 or SPF colonized pups weaned onto FR or FF diets. Bold horizontal bar represents the mean and lighter error bars the S.E.M. (n=5–12). In panels D, P, two-way ANOVA and post hoc test with Original FDR method of Benjamini and Hochberg was used for statistics. Related to main text **Figure 1**.

**Supplemental Figure 2.**
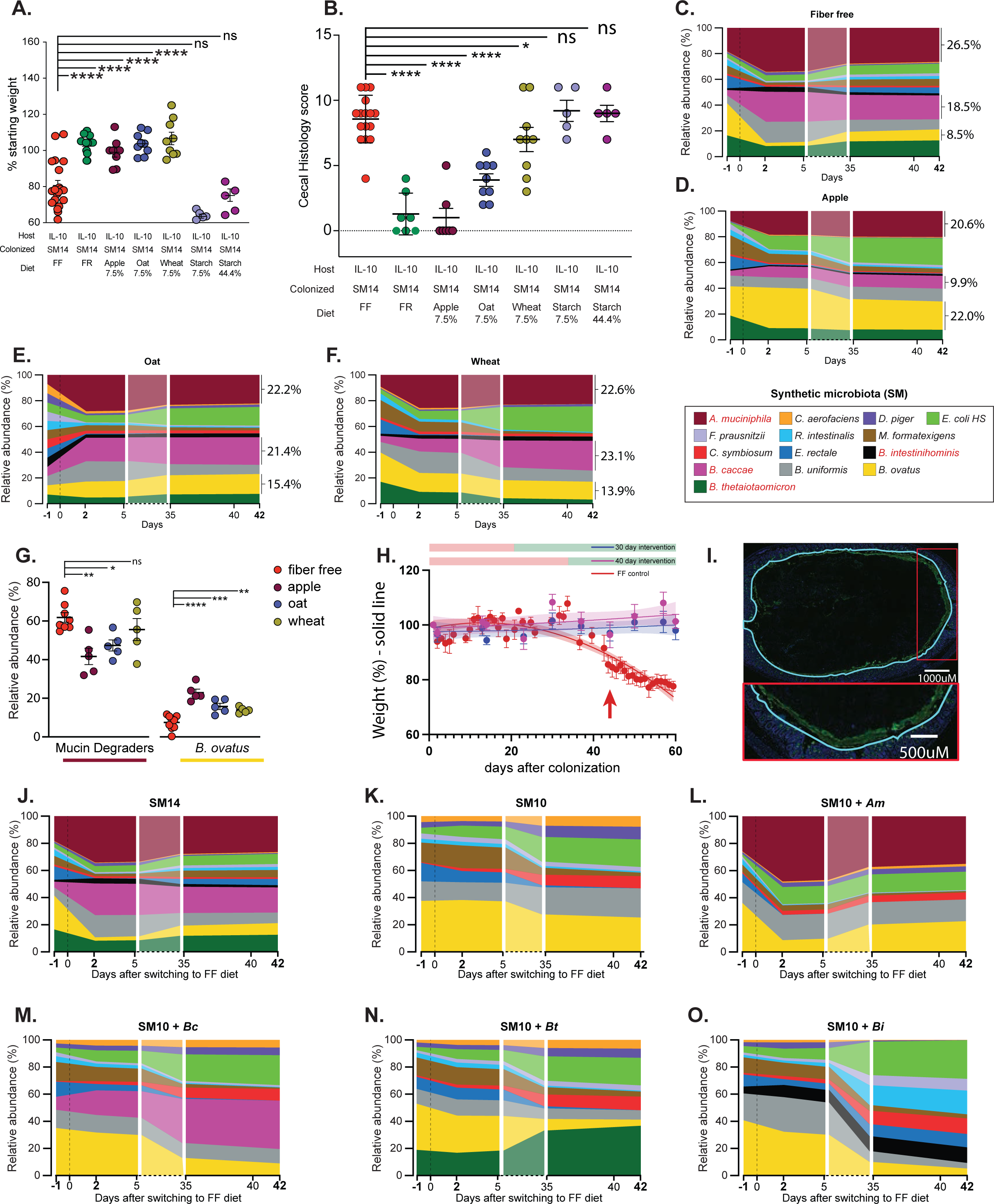
Fiber supplementation reduces inflammation severity. **A.** Endpoint weights of mice fed variations of the FF diet with glucose replaced by fiber from apple, oat or wheat, or soluble starch. Fiber or starch were added at the concentrations listed below each treatment and an equal amount of glucose was omitted. Bold horizontal bar represents the mean and lighter error bars the S.E.M. (n=5–20). **B.** Cecal histology scores of the treatment groups shown in A. (n=5–20). **C.–F.** Relative abundance plots of the SM14 bacteria in *Il10^-/-^* mice fed fiber-enriched versions of the FF diet and containing either no added fiber (C.) or purified fiber from apple (D.), oat (E.) or wheat (F.). Relative abundance values for *A. muciniphila*, *B. caccae* and *B. ovatus* are shown next to each plot. **G.** Relative abundance comparison of all 4 mucin degraders compared to the fiber-degrader *B. ovatus* in adult mice colonized with SM14 and fed FF or the fiber supplemented diets. Note that the apple diet which had the lowest inflammation had the largest increase in *B. ovatus* and the largest decrease in mucin degrading bacteria (n=5– 8). **H.** Weight trajectories of the mice that were returned to the FR diet at either 30d or 40d. **I.** Representative mucus staining for SM10 colonized mice used to measure thickness with Bacspace software. α-Muc2: green, DAPI: blue, prediction of epithelium/mucus interface: teal. **J.-O.** Relative abundance plots of the SM14 bacterial members in *Il10^-/-^* mice with all bacteria present (J.), SM10 (K.), SM10 + *A. muciniphila* (L.), SM10 + *B. caccae* (M.), SM10 + *B. thetaiotaomicron* (N.), SM10 + *B. intestinihominis* (O.). In panels A, B, G, two-way ANOVA and post hoc test with Original FDR method of Benjamini and Hochberg was used for statistics. Related to main text **Figures 1 and 2**.

**Supplemental Figure 3.**
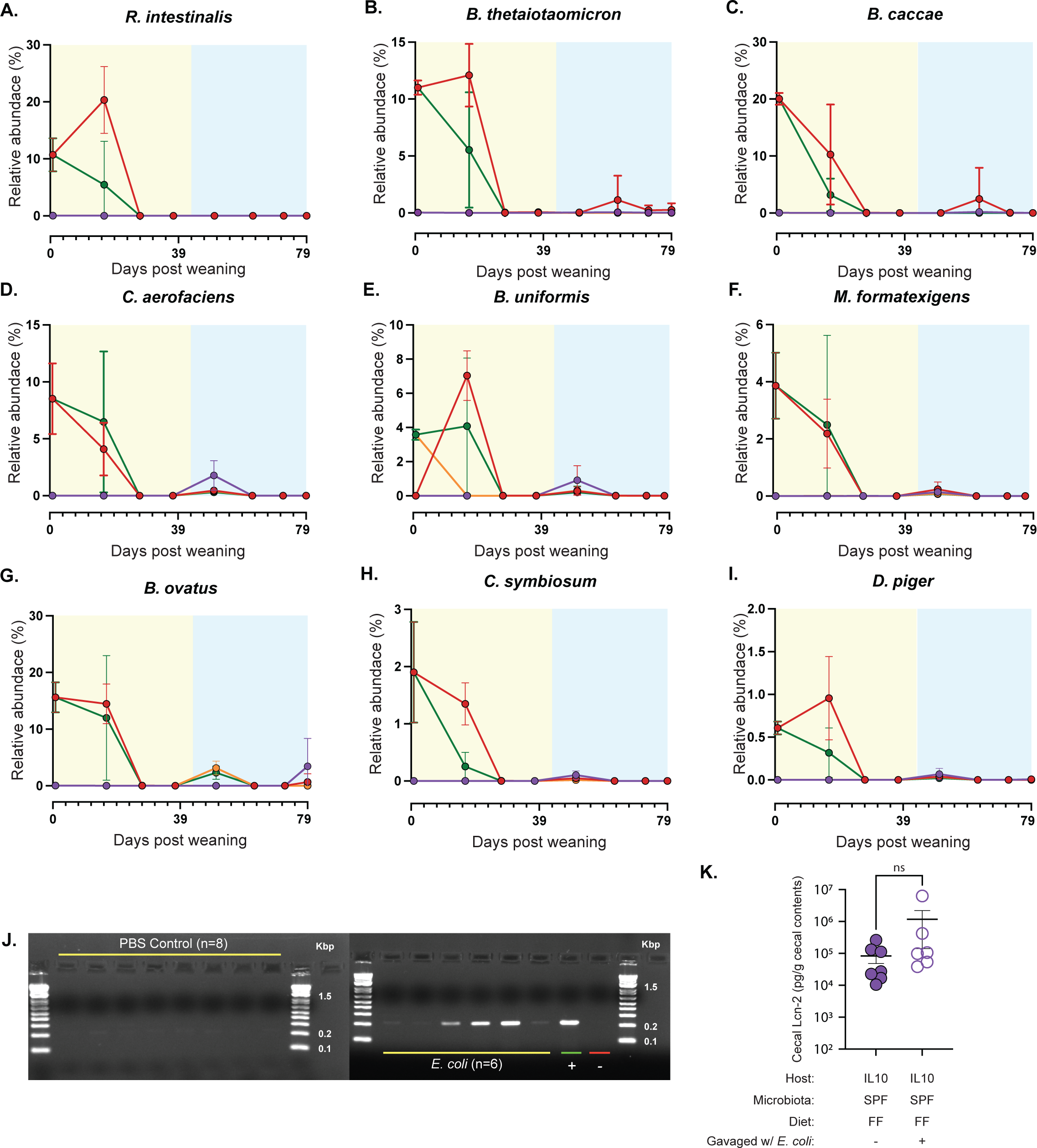
Effects of co-housing SM14 and SPF mice fed FR and FF diets. **A.-I.** 16S rRNA gene based measurements of the remaining SM14 bacteria in co-housed mice: *R. intestinalis* (**A.**)*, B. thetaiotaomicron* (**B.**)*, B. caccae* (**C.**), *C. aerofaciens* (**D.**), *B. uniformis* (**E.**), *M. formatexigens* (**F.**), *B. ovatus* (***G.***), *C. symbiosum* (**H.**) and *D. piger* (**I.**). **J.** PCR analysis using *E. coli* HS specific primers^13^ for the presence of *E. coli* HS in mice that were mock gavaged weekly with PBS (left) or *E. coli* HS (right). **K.** Cecal LCN2 measurements in SPF mice that were gavaged weekly with *E. coli* HS and fed the FF diet. (n=6-7, one-tailed Student’s t-test and Wilcoxon test). Related to main text **Figure 3**.

**Supplemental Figure 4.**
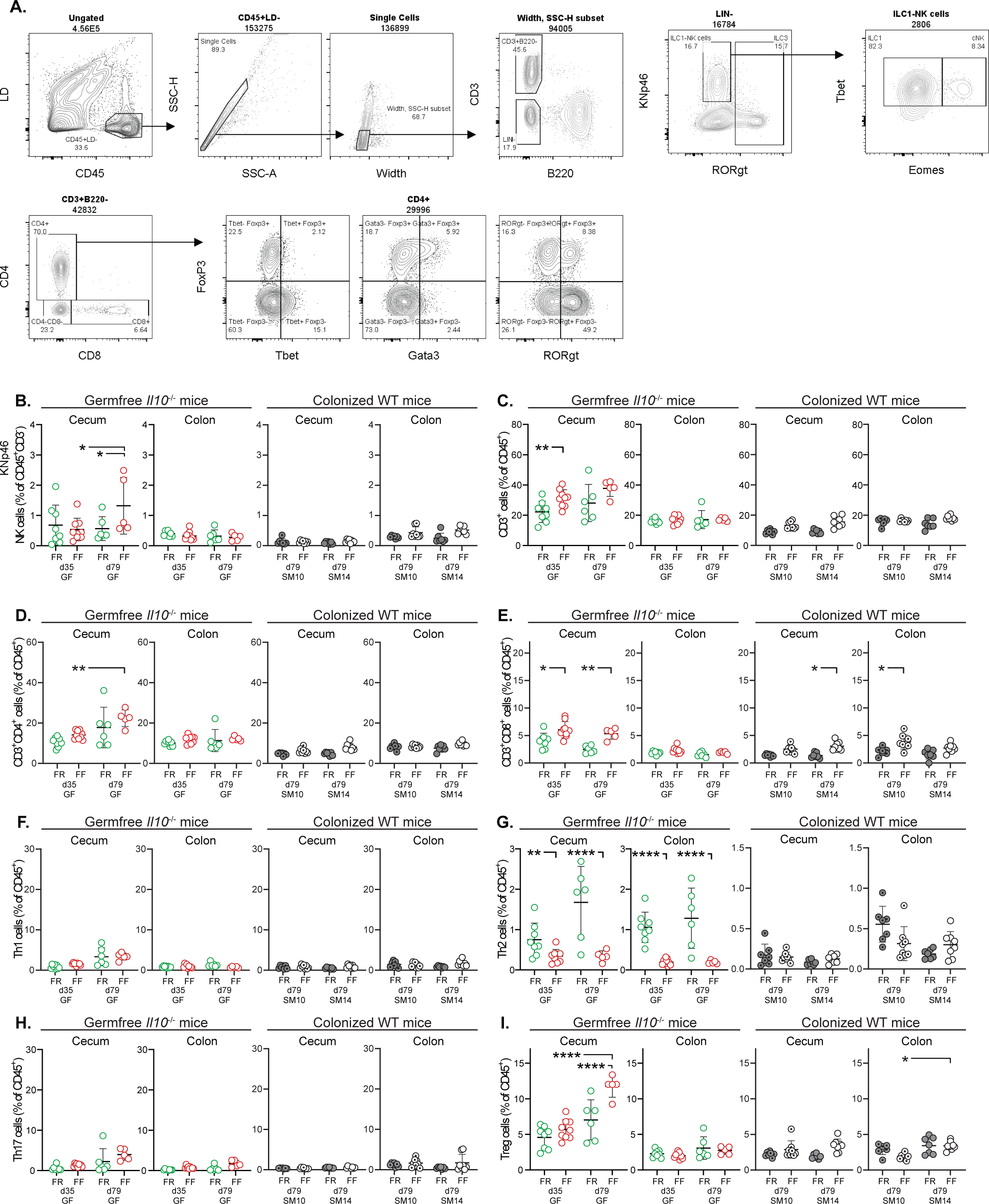
Fiber deprivation does not cause colitis in the absence of genetic predisposition (WT mice) or microbiota (germfree). **A.** Gating Strategy for the analysis of T cells and NK cells in the cecal and colonic lamina propria. **B.-I.** Proportions of immune cell populations in the cecum and colon of GF *Il10^-/-^* mice and SM14- or SM10-colonized WT mice in the cell types listed on the y-axis of each panel (n=5**–**9, two-way ANOVA and post hoc test with Original FDR method of Benjamini and Hochberg). Data are represented as mean ± SD and. For each cell population, data were analyzed along with data from Fig. 3 and S5. *p < 0.05; **p < 0.01; ***p < 0.001; ****p < 0.0001. Related to main text **Figure 5**.

**Supplemental Figure 5.**
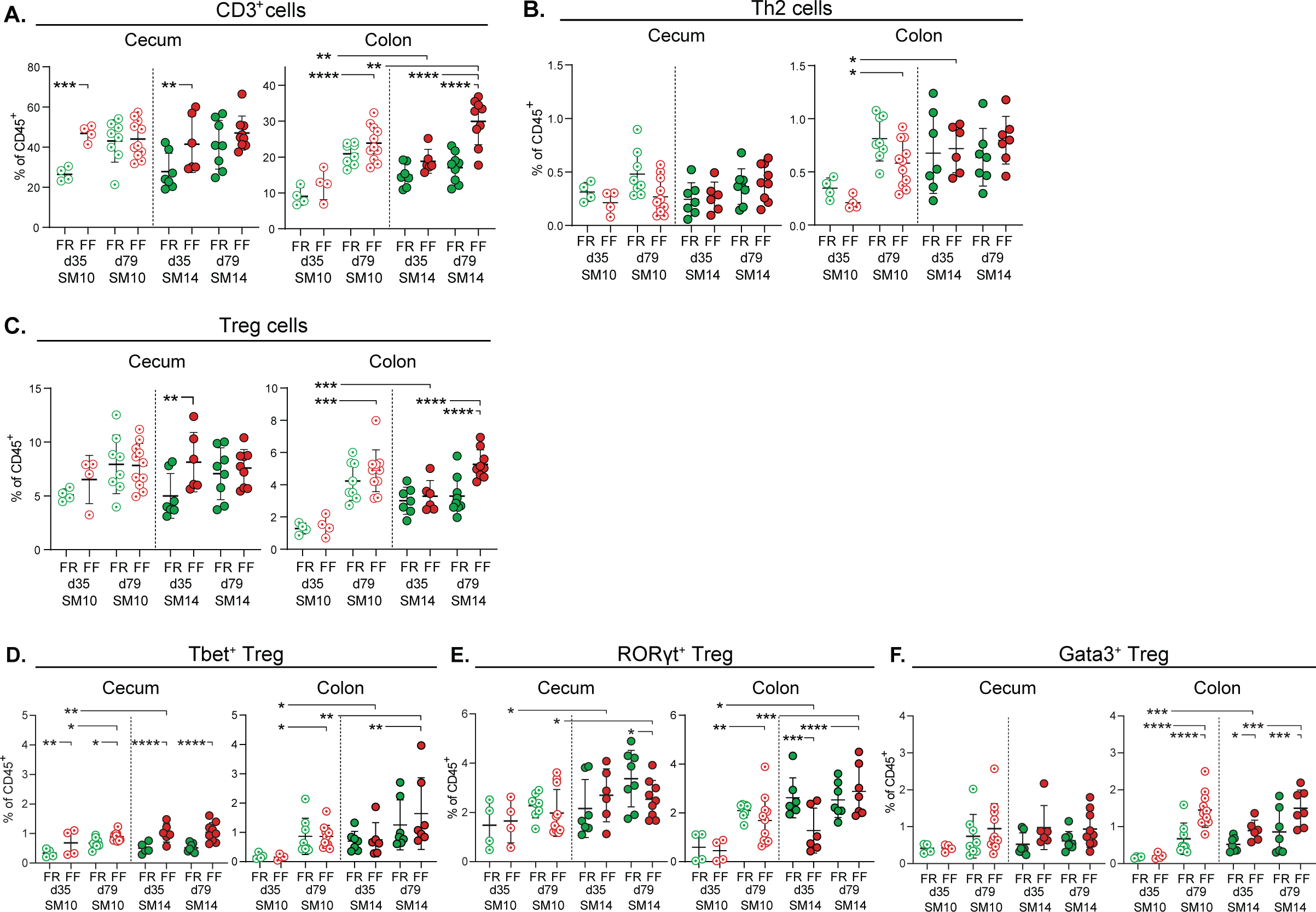
Mucin-degrading bacteria promote Th1 immune responses in a diet-dependent manner. **A–F.** Proportion of CD3^+^ (**A.**), Th2 (**B.**), Treg cells (**C.**), Tbet^+^ Treg (**D.**), RORyt^+^ Treg (**E.**) and Gata3^+^ Treg (**F.**) cells among CD45^+^ cells in the cecum and colon of SM10- and SM14-colonized *Il10^-/-^*mice (n=4**–**11). Data are represented as mean ± SD. *p < 0.05; **p < 0.01; ***p < 0.001; ****p < 0.0001. In panels B**–**G, two-way ANOVA and post hoc test with Original FDR method of Benjamini and Hochberg was used for statistics. Related to main text **Figures 5-6**.

**Supplemental Figure 6.**
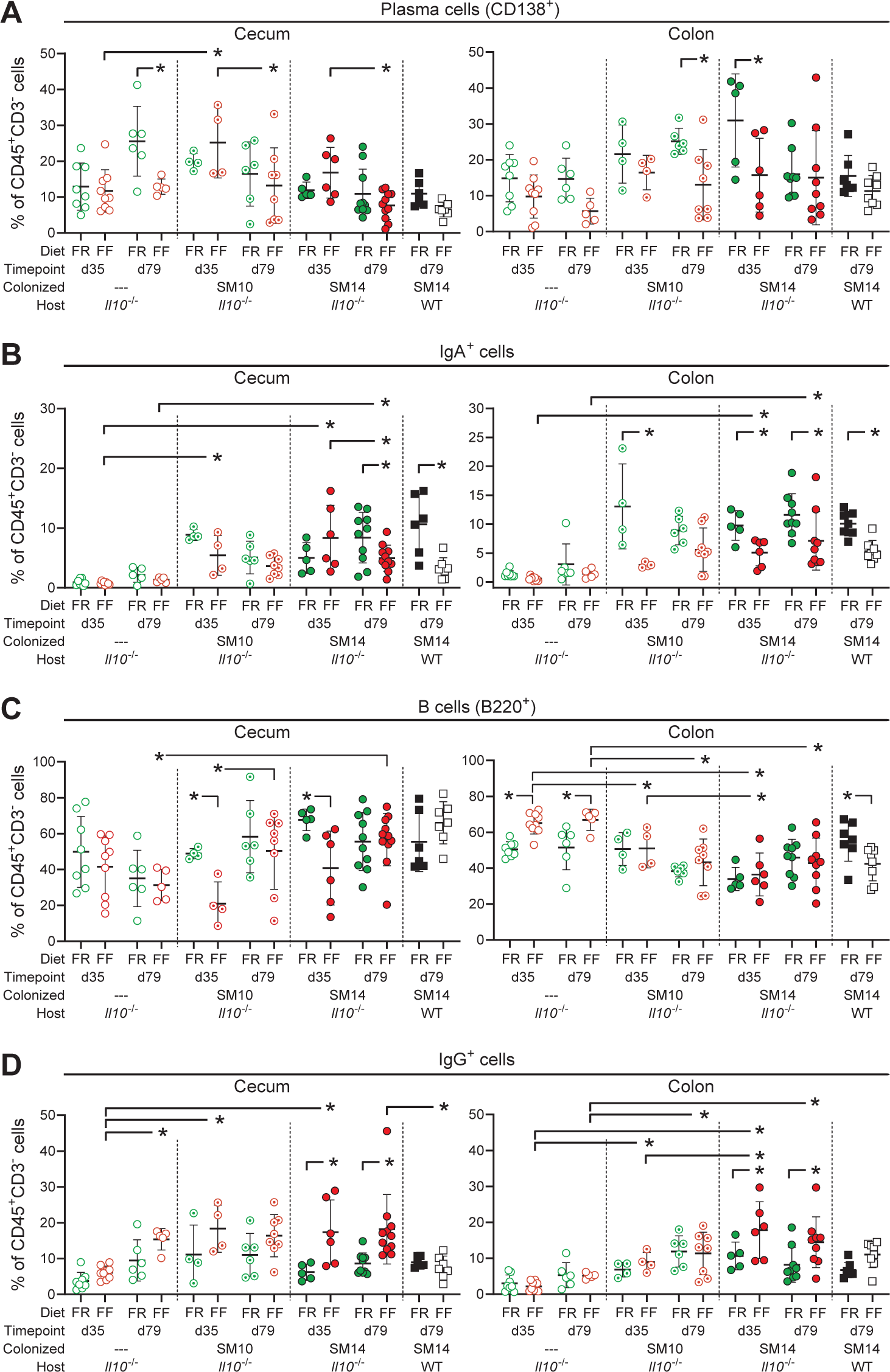
Fiber deprivation alters Ig–producing cell populations. **A.–D.** Proportion of plasma cells (**A.**), IgA-producing cells (**B.**), B cells (**C.**) and IgG-producing cells (**D.**) among CD3^-^CD45^+^ cells in the cecum and colon of GF, SM10- and SM14-colonized *Il10^-/-^* and WT mice (n=4**–**11, two-way ANOVA and post hoc test with Original FDR method of Benjamini and Hochberg). Related to main text **Figure 7**.

**Supplemental Figure 7.**
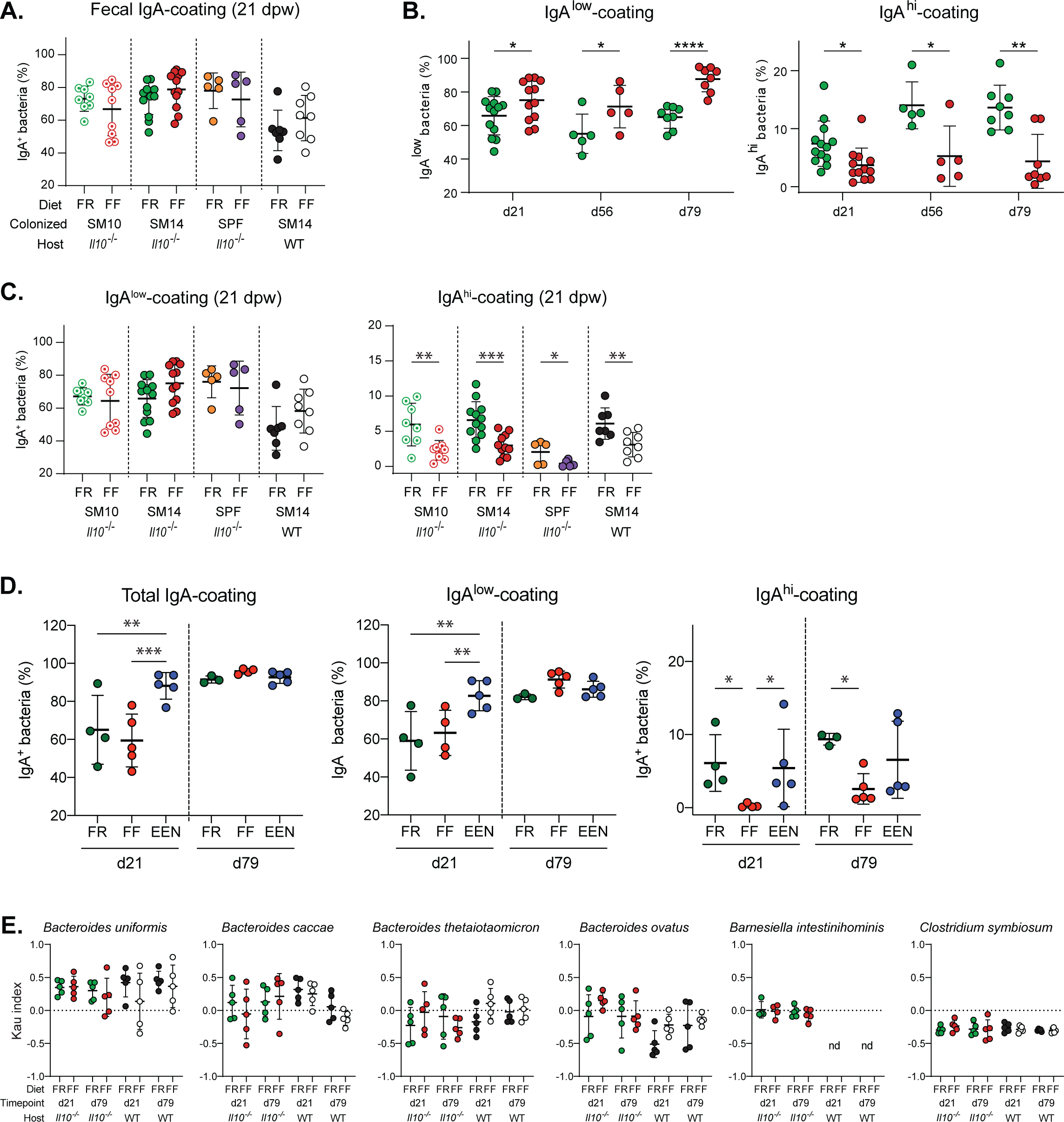
Fiber deprivation alters IgA–bacteria interactions. **A.** Percentages of total IgA-coated bacteria as shown in the feces of SM10-, SM14- or SPF-colonized *Il10^-/-^* mice and SM14-colonized WT mice fed the FR or the FF diet for 21 days (n=5**–**8, two-way ANOVA and post hoc test with the two-stage linear step-up procedure of Benjamini, Krieger and Yekutieli). **B–C.** Percentages of IgA^low^- and IgA^high^-coated bacteria in the feces SM14-colonized *Il10^-/-^* mice over time (B.) or indicated groups at 21 dpw (C.) (n=5**–**8, two-way ANOVA and post hoc test with the two-stage linear step-up procedure of Benjamini, Krieger and Yekutieli). **D.** Percentages of total (left), IgA^low^ (middle) and IgA^high^ (right) -coated bacteria in the feces of SM14-colonized *Il10^-/-^* mice fed the FR, FF or EEN diet for 21 or 79 days (n=3–5, two-way ANOVA and post hoc test with the two-stage linear step-up procedure of Benjamini, Krieger and Yekutieli). **E.** IgA-coating index (Kau index) of fecal bacteria from SM14-colonized *Il10^-/-^* and WT mice (n=2–5, multiple unpaired t-test and post hoc test with the two-stage linear step-up procedure of Benjamini, Krieger and Yekutieli). Data are represented as mean ± SD. *p < 0.05; **p < 0.01; ***p < 0.001; ****p < 0.0001. Related to main text **Figure 7**.

## STAR METHODS

### Key resources table

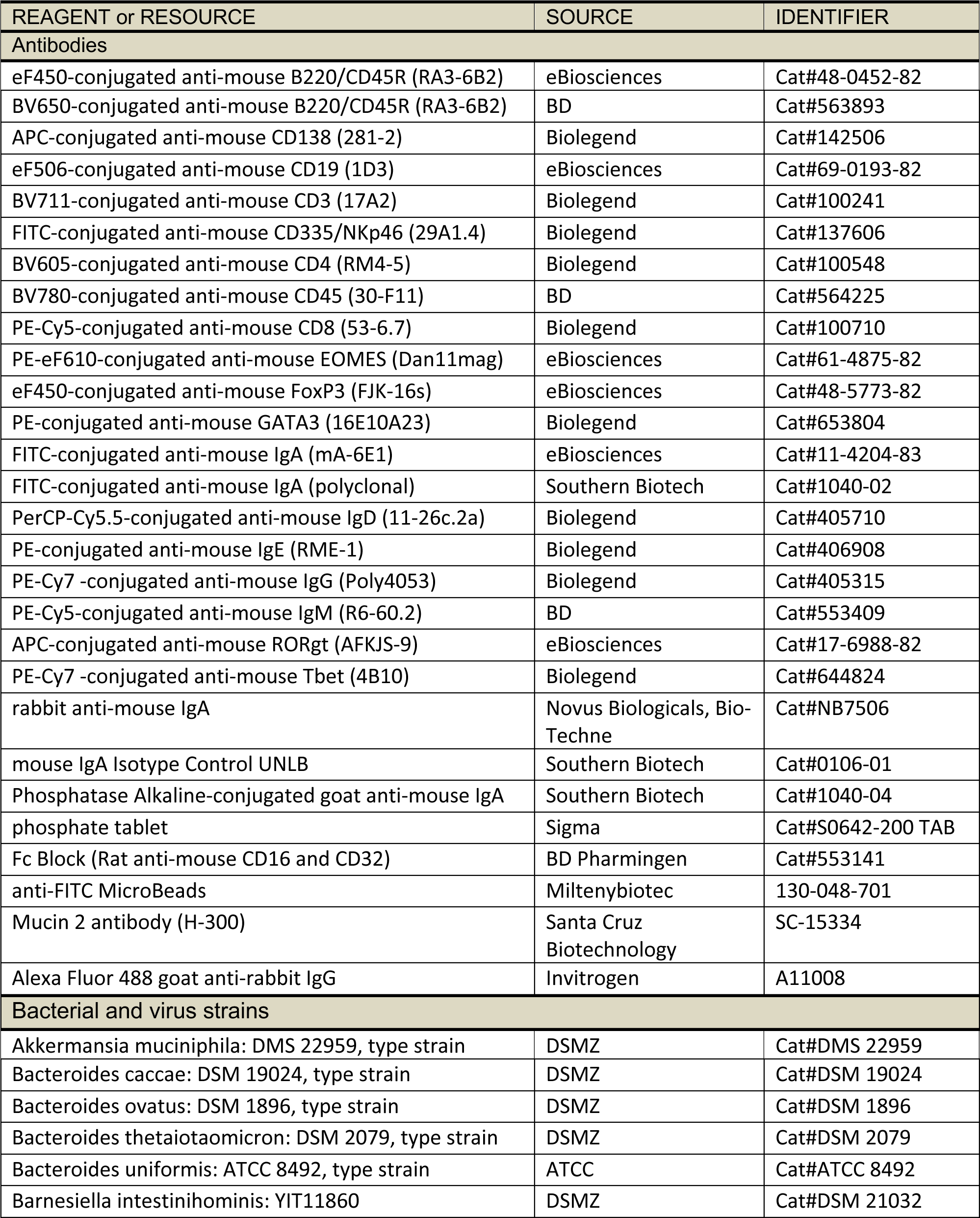
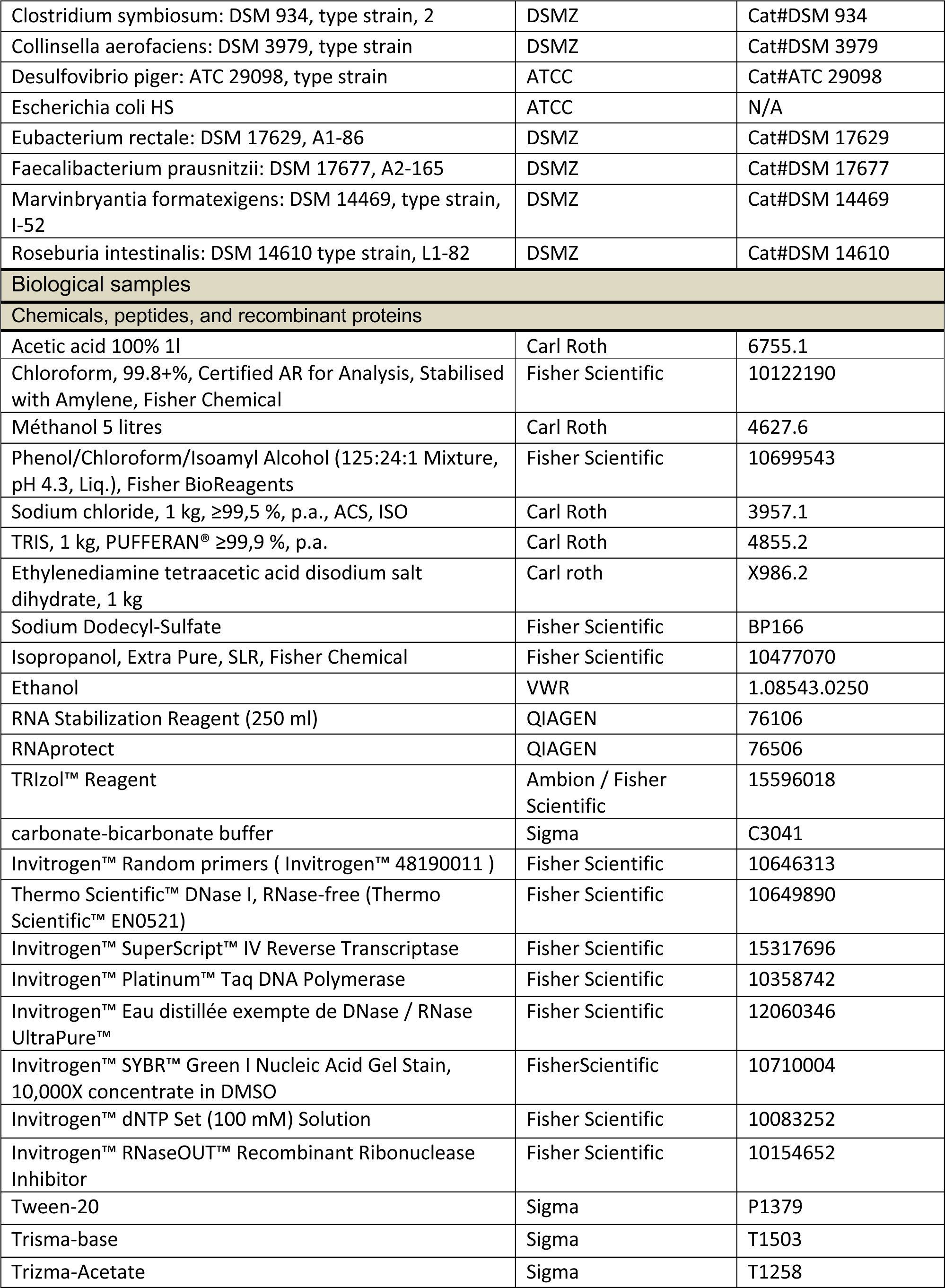
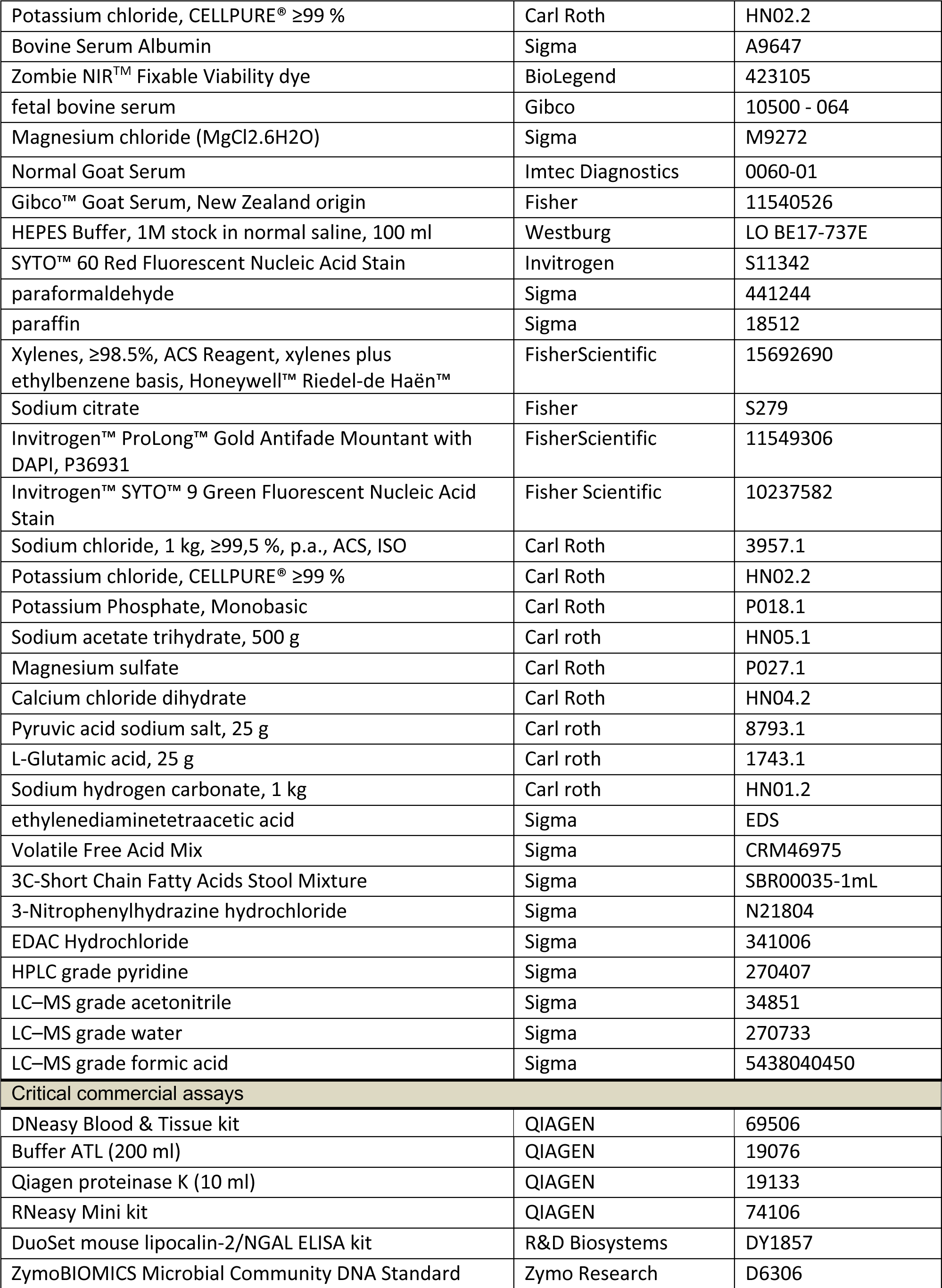
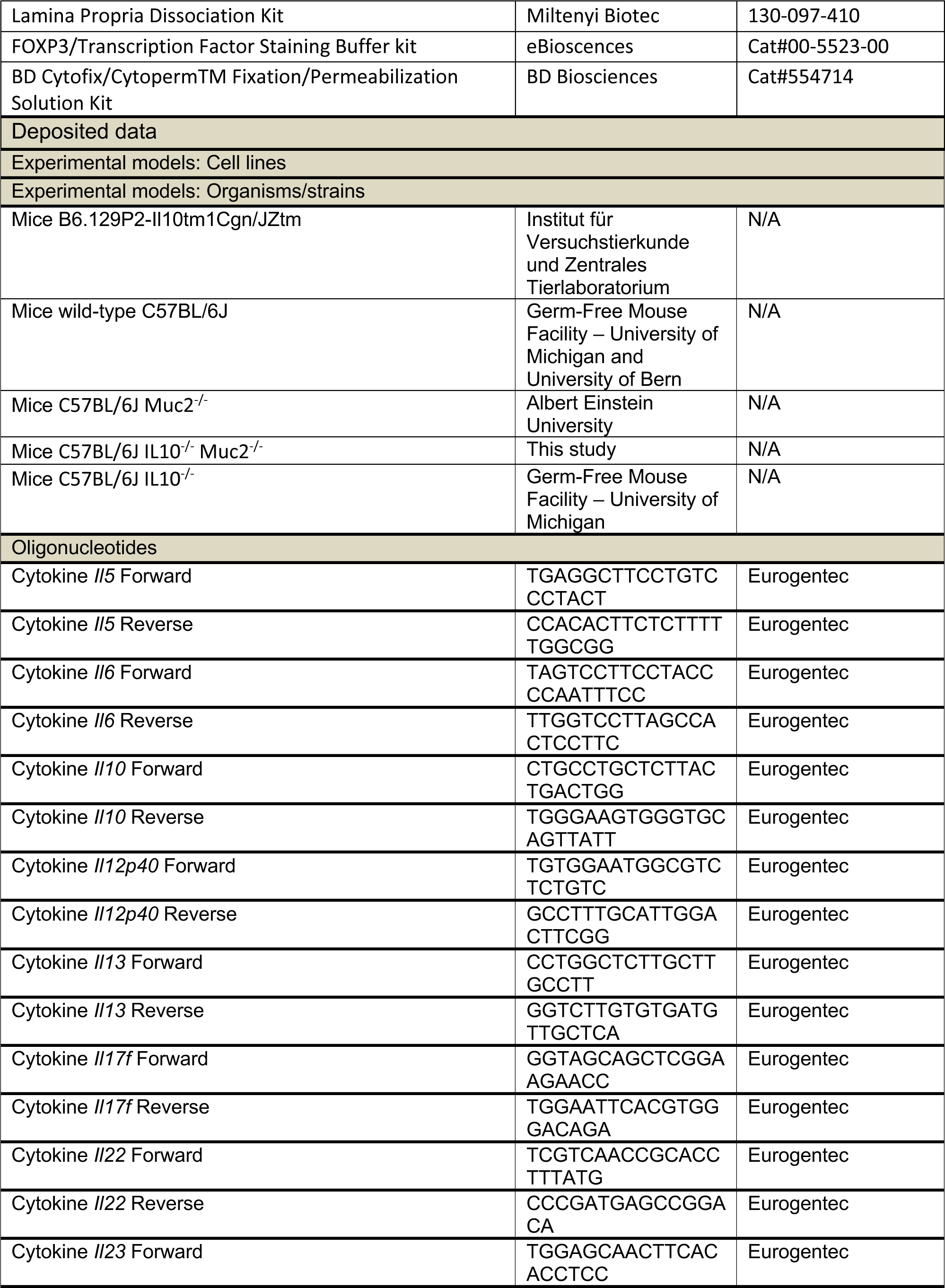
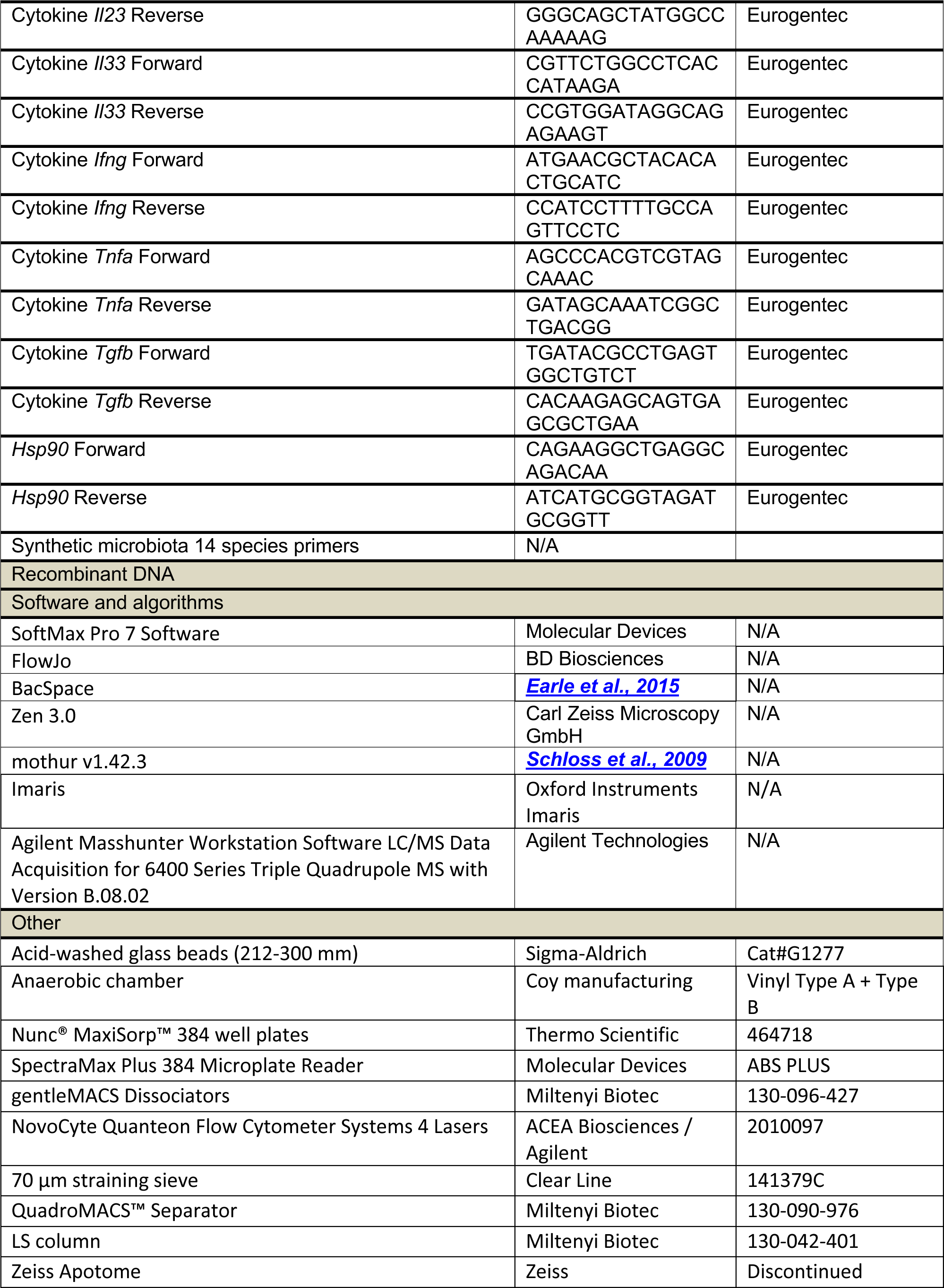
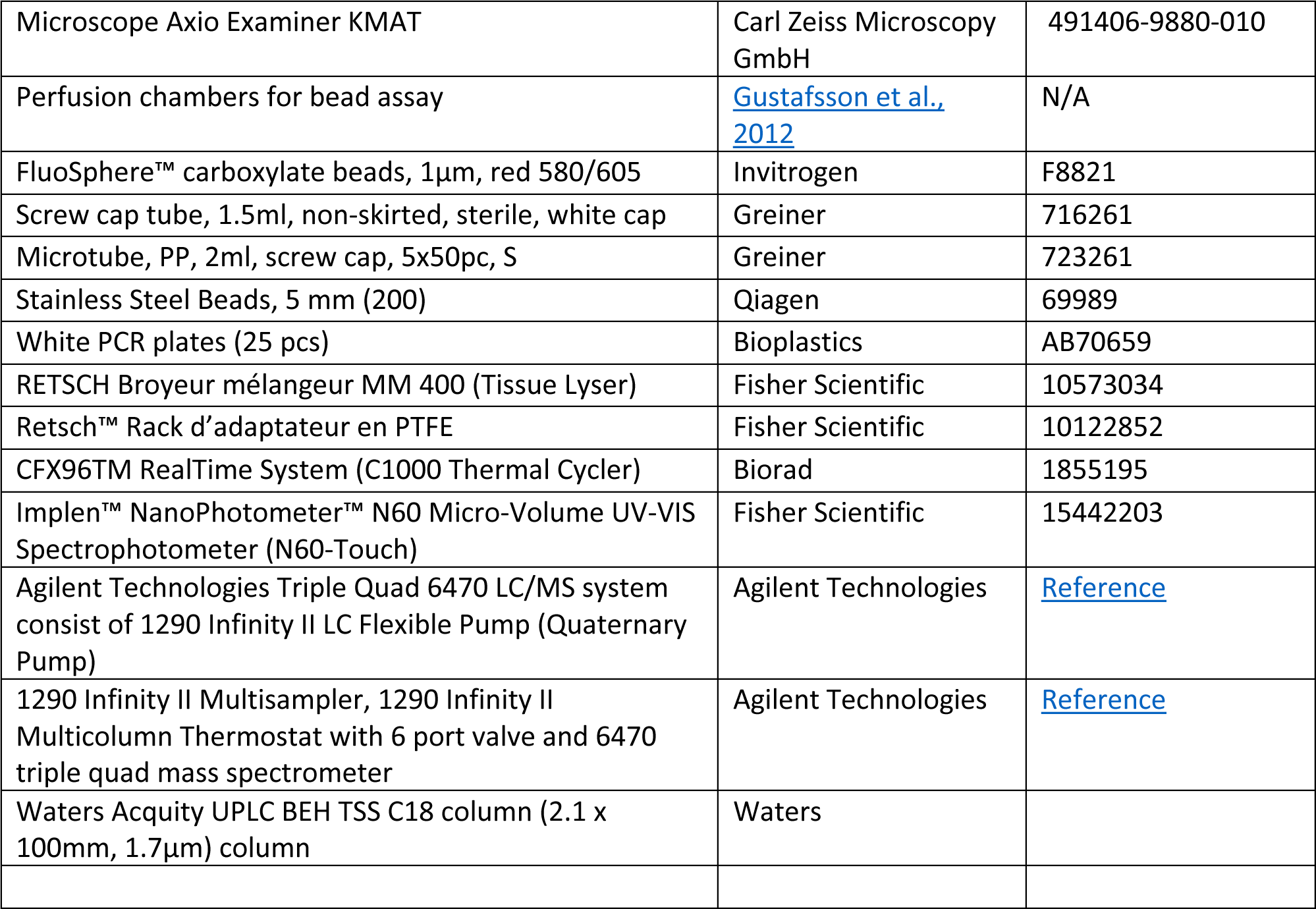

## LEAD CONTACT AND MATERIALS AVAILABILITY

Bacterial strains and data associated with this study are available to interested parties upon request to.

## EXPERIMENTAL MODEL AND SUBJECT DETAILS

### Animal models, colonization diet and sample processing

Animal experiments at the University of Michigan followed protocols approved by the University of Michigan Institutional Animal Care and Use Committee (IACUC) and were supervised by licensed verterinarians. Germ-free B6.129P2-Il10tm1Cgn/JZtm mice (Institut für Versuchstierkunde und Zentrales Tierlaboratorium, Hannover, Germany) and wild-type C57BL/6J (University of Bern, Switzerland) were bred at the animal facility of the University of Luxembourg and underwent protocols approved by the Animal Experimentation Ethics Committee of the University of Luxembourg and the Ministre de l’Agriculture, de la Viticulture et du Development rural du Grand-Duché du Luxembourg (LUPA 2020/20). Germfree males and females were colonized at 7–10 weeks of age and none of these mice was involved in any previous experiments/treatments. Mice were either housed alone or in groups of appropriate gender, litter, and dietary experiments. Food and autoclaved distilled water were provided ad libitum. Mice were weighed and monitored at least weekly for diarrhea, prolapses, and general health state and this was increased to daily monitoring in animals that began experiencing weight loss. The *Muc2*^-/-^ mice, crossed to C57BL/6J background, were obtained from Leonard Augenlicht (Albert Einstein University). For the generation of *Il10*^-/-^ *Muc2*^-/-^ double knockout (DKO), the single KO mice were crossed to obtain double heterozygous mice until generation of *Il10*^-/-^ *Muc2*^+/-^. Full double KO mice developed spontaneous prolapse in germ-free condition during prolonged breeding periods, thus, the *Il10*^-/-^ *Muc2*^+/-^ mice were used to generate the double KO mice used for all experiments. Genotyping was performed by extracting DNA from ear punches (Transnetyx). Mice were gavaged with the synthetic microbiota at 6–8 weeks and proceeded to diet change as previously described. The synthetic microbiota bacteria were grown in their respective medium^13^ or a modified YCFA medium^43^ prior to community assembly for gavages. The bacteria were cultivated under anaerobic atmosphere maintained with a gas mixture (85% N_2_, 10% H_2_, 5% CO_2_) to an optical density (absorbance 600nm) ranging from 0.5 to 1.0. The communities were assembled by mixing equal volumes of each specific bacterium and aliquoted into sealed screw cap tubes with its own headspace. Each mouse was gavaged with 0.2 mL of its specific community, depending on the experiment, with freshly prepared inocula for two-three consecutive days. A humane endpoint was used for mice that lost ≥20% of their starting weight and these mice were counted as lethalities in the weight loss and survival curves shown. Animals were euthanized using CO_2_ asphyxiation for 5 minutes followed by cervical dislocation. The gastrointestinal tracts were retrieved quickly, to prevent autolysis, and the sections separated. Cecal and colon contents of each animal were flash frozen in liquid nitrogen and kept at -80°C until further use.

The FR diet (Lab Diet 5013) and FF diet (Envigo-Teklad TD.130343) have been previously described^13^. The EEN diet employed was Nestle Nutren 1.5, which was lyophilized and sent to Envigo rodent diets to be formed into pellets, bagged and gamma irradiated prior to use. Apple (Vitacel AF401), oat (Vitacel HF600-30) and wheat (Vitacel WF200) fibers were from J. Rettenmaier (Schoolcraft, MI, USA) and added to the FF diet at 7.5% w/w with a corresponding decrease in the amount of glucose in the FF diet. Soluble, highly digestible starch was provided by Cargill (Gel Instant 12412).

## METHOD DETAILS

### DNA extraction

DNA from fecal and cecal samples were isolated using a bead-beating phenol:chloroform extraction method followed by DNeasy Blood & Tissue Kit (QIAGEN, USA). In short, samples were weighed between 10-50mg and combined with acid-washed glass beads (212-300mm; Sigma-Aldrich, USA), 500uL Buffer A (200 mM NaCl, 200 mM Tris, 20 mM EDTA, 210 uL SDS (20% w/v, filter-sterelized), and 500 uL phenol:chloroform (Thermo Fisher Scientific, USA). The samples were disrupted using a Mini-Beadbeater-16 (Biospec Products, USA) for 3 minutes at room temperature and centrifuged (10,000 rpm, 4°C, 3 minutes). The aqueous phase was recovered and mixed with an equal volume of phenol:chloroform by gentle inversion and centrifuged (10,000 rpm, 4°C, 3 minutes). The remaining aqueous phase was recovered and mixed with 500 µL of chloroform, mixed by gentle inversion, and centrifuged (10,000 rpm, 4°C, 3 minutes). Recovered aqueous phase was mixed with 1 volume of isopropanol and 1/10 volume of 3M sodium acetate, and stored at -80°C for 20 minutes for DNA precipitation. Samples were centrifuged (15000 rpm, 4°C, 20 minutes), supernatant discarded, washed with 70% ethanol, air-dried, and resuspended in nuclease-free water. The sample DNA extracts were further purified with DNeasy Blood & Tissue kit, following manufacturer protocol (QIAGEN).

### RNA extraction, reverse transcription and qPCR

Freshly retrieved ileal, cecal and distal colon tissues were transferred into RNAlater^TM^ (QIAGEN) and kept at 4°C up to a week. Then, RNAlater^TM^ was removed and tissues were stored at -80°C until further use. Frozen tissues were transferred into 1 mL of TRIzol reagent (Invitrogen^TM^), homogeneized with a 5 mm metal bead on a bead beater for 8 min at 30Hz and centrifuged for 3 min at 13000 rpm, 4°C. The supernatant was recovered, mixed thoroughly with 200 µl of chloroform and incubated at room temperature for 2-3 min before a centrifigation for 15 min at 13000 rpm, 4°C. The aqueous phase was recovered, mixed again with an equal amount of chloroform and centrifuged for 15 min at 13000 rpm, 4°C. The aqueous phase was recovered, mixed by inversion with 500 µl of isopropanol, incubated for 10 min at room temperature, and centrifuged for 10 min at 13000 rpm, 4°C. The pellet was washed with 1 ml Ethanol 70% and centrifuged for 5 min at 10000 rpm. The supernatant was discarded and the pellet dried for 5-10 min at 37°C, resuspended with 50 µl nuclease-free water and incubated for 15 min at 56°C. Finally, samples were treated with DNase following the Thermo Scientific DNase1, RNase-free Protocol, and RNA were purified with the RNeasy Mini kit (QIAGEN) according to manufacturer instructions. Final RNA concentrations were quantified by Nanodrop.

### Lipocalin-2 (LCN2) measurements in cecal and fecal contents

Frozen cecal and fecal samples were used to quantify the presence of lipocalin-2 protein (LCN2) by enzyme linked immunosorbent assay (ELISA). Samples previously frozen (cecal, - 80°C, and fecal, -20°C) were weighed in new tubes between 5-5 mg. Samples were kept over dry ice during the weighing. Samples were resuspended in 1 mL PBS (pH 7.4), vortexed for 30s, and kept at 4°C overnight to homogenize. Samples were then extensively vortexed to a homogeneous solution. To measure the LCN2 levels, a DuoSet mouse lipocalin-2/NGAL ELISA kit (R&D Biosystems, USA) was employed using several dilutions of sample homogeneous solution. Quantification was done following manufacturer protocol.

### Soluble IgA measurements in cecal and fecal contents

To measure soluble IgA levels, Nunc® MaxiSorp™ 384 well plates (Sigma-Aldrich) were coated overnight with 10 ng/well rabbit anti-mouse IgA (Novus Biologicals, Bio-Techne NB7506) in 20 µl/well of carbonate-bicarbonate buffer (Sigma, Ref.: C3041). After four washes with Washing Buffer (1% Tween-20, 154mM Sodium Chloride and 10mM Trisma-base), 75μl of Blocking Buffer (15 mM Trizma-Acetate, 136 mM Sodium Chloride, 2 mM Potassium Chloride and 1% (w/v) BSA (Bovine Serum Albumin)). After 2h at room temperature, wells were washed again. Sample homogeneous solution and standards (mouse IgA Isotype Control UNLB, Southern Biotech, Imtec Diagnostics, Ref: 0106-01) were diluted in Dilution Buffer (15 mM Trizma-Acetate, 136 mM Sodium Chloride, 2 mM Potassium Chloride, 0.1% (w/v) Tween-20, and 1% BSA) and incubated into the plate at 20 µl/well, room temperature for 90 min. After washing, 20 µl/well of a Phosphatase Alkaline-conjugated goat anti-mouse IgA (Southern Biotech, Imtec diagnostics, Ref: 1040-04), diluted 1/1000 in Dilution Buffer, was added and incubated at room temperature for 90 min. After a final wash, 40 µl/well of substrate (1 phosphate tablet (Sigma, ref S0642-200 TAB) dissolved in 10 mL Substrate Buffer (1 mM 2-Amino-2-methyle-1-propanole, 0.1 mM MgCl2.6H2O)) was added. The plate was incubated at 37°C for 60 min before the absorbance was measured at 405 nm using an ELISA plate reader (SpectraMax Plus 384 Microplate Reader from Molecular Devices; Software: SoftMax Pro 7 Software, Molecular Devices). The IgA concentration was determined for each sample using the formulated standard curve.

### Short- and branched-chain fatty acid quantification

Short-chain and branched-chain fatty acids (SCFAs) standards mixture was obtained from Sigma (CRM46975). 13C-short chain fatty acid stool mixture (Sigma, SBR00035-1mL) was used as the internal standard (IS). Analytical reagent-grade 3-nitrophenylhydrazine (3NPH)·HCl (Cat#N21804), EDAC·HCl (Cat#341006); HPLC grade pyridine (Cat#270407); LC–MS grade acetonitrile (Cat#34851), water (Cat#270733), and formic acid (Cat#5438040450) were also purchased from Sigma–Aldrich. The working standard solutions were created by performing serial dilution from the 10mM stock solution down to nM range using freshly prepared 50% (v/v) aqueous acetonitrile in water. The chemical derivatization protocol was modified from Han et al. ^44^. Briefly, 20µL of the working standard solutions or samples was mixed with 40µL of 200mM 3NPH in 50% aqueous acetonitrile, 120mM EDAC- 6% (v/v) pyridine solution in the same solvent and 4µL of the IS in a Verex glass vial. The mixture was reacted at 40°C for 30 min. After reaction, 96µL of 0.1% formic acid in 10% acetonitrile solution was added to the mixture to quench the reaction. 30µL of the reaction solution was then transferred to a new HPLC vial and 2-µL aliquot of each solution was injected into the LC-MS/MS instrument. Each modified SCFA was optimized in Agilent MS for detection through Agilent Optimizer 2.0. All optimized SCFAs information was combined, and a LC-MRM MS method was created. Retention time for each SCFA was determined from two transitions. Then the MRM MS method was transformed into a dynamic MRM MS or dMRM MS method with all the RTs and MS information for the final LC-MS/MS acquisition method.

LC-MS/MS analysis was performed on the Agilent Technologies Triple Quad 6470 LC/MS system consist of 1290 Infinity II LC Flexible Pump (Quaternary Pump), 1290 Infinity II Multisampler, 1290 Infinity II Multicolumn Thermostat with 6 port valve and 6470 triple quad mass spectrometer. Agilent Masshunter Workstation Software LC/MS Data Acquisition for 6400 Series Triple Quadrupole MS with Version B.08.02 is used for calibration, compound optimization and sample data acquisition.

A Waters Acquity UPLC BEH TSS C18 column (2.1 x 100mm, 1.7µm) column was used with mobile phase A) consisting of 0.1% formic acid in water; mobile phase (B) consisting of 0.1% formic acid in acetonitrile. Gradient program: mobile phase (B) was held at 15% for 1 min, increased to 55% in 19 min, then to 99% in 20 min and held for 2 min before going to initial condition and held for 4 min. The column was at 40 °C and 2 µl of sample was injected into the LC-MS with a flow rate of 0.3 ml/min. Calibration of the 6470 MS was achieved through Agilent ESI-Low Concentration Tuning Mix. Source parameters: Gas temp 300 °C, Gas flow 5 l/min, Nebulizer 45 psi, Sheath gas temp 250 °C, Sheath gas flow 11 l/min, Capillary - 3500 V, Delta EMV -200 V. Dynamic MRM scan type is used with 0.07 min peak width. dMRM transitions and other parameters for each compound were list in a separate sheet. Delta retention time of plus and minus 1 min, fragmentor of 40 eV and cell accelerator of 5 eV were incorporated in the method. Data analysis was performed by Agilent Mass Hunter Quantitative Analysis for QQQ B.10.00 for integration. Results were exported to CVS file for further analysis.

### Lamina propria cell extraction and flow cytometry analysis

Cecal and colonic lamina propria cells were extracted using the Lamina Propria Dissociation Kit and gentleMACS Dissociators (Miltenyi Biotec, Germany) according to the manufacturer’s instruction. After digestion, cells were resuspended in PB buffer (PBS, pH 7.2, 0.5 % BSA) and counted. For analysis of T cells and NK cells with the expression of transcription factors, the FOXP3/Transcription Factor Staining Buffer kit (eBioscences – 00-5523-00) was used along with the following anti-mouse antibodies: BV605-conjugated anti-CD4 (Biolegend, RM4-5; 1/700), BV650-conjugated anti-B220 (BD Biosciences, RA3-6B2; 1/88), BV711-conjugated anti-CD3 (Biolegend, 17A2; 1/88), BV780-conjugated anti-CD45 (BD Biosciences, 30-F11; 1/88), FITC-conjugated anti-CD335/NKp46 (Biolegend, 29A1.4; 1/100), PE-Cy5-conjugated anti-CD8 (Biolegend, 53-6.7; 1/700), eF450-conjugated anti-FoxP3 (eBiosciences, FJK-16s; 1/200), PE-conjugated anti-GATA3 (Biolegend, 16E10A23; 1/44), PE-eF610-conjugated anti-EOMES (eBiosciences, Dan11mag; 1/100), PE-Cy7 -conjugated anti-Tbet (Biolegend, 4B10; 1/44), APC-conjugated anti-RORgt (eBiosciences, AFKJS-9; 1/22). For B cells analysis and immunoglobulin expression, the BD Cytofix/Cytoperm^TM^ Fixation/Permeabilization Solution Kit (BD Biosciences – 554714) was used along with the following anti-mouse antibodies: eF450-conjugated anti-B220/CD45R (eBiosciences, RA3-6B2; 1/700), eF506-conjugated anti-CD19 (eBiosciences, 1D3; 1/88), BV711-conjugated anti-CD3 (Biolegend, 17A2; 1/88), BV780-conjugated anti-CD45 (BD Biosciences, 30-F11; 1/88), APC-conjugated anti-CD138 (Biolegend, 281-2; 1/100), FITC-conjugated anti-IgA (eBiosciences, mA-6E1; 1/700), PerCP-Cy5.5-conjugated anti-IgD (Biolegend, 11-26c.2a; 1/200), PE-conjugated anti-IgE (Biolegend, RME-1; 1/44), PE-Cy5-conjugated anti-IgM (BD Biosciences, R6-60.2; 1/100), PE-Cy7 -conjugated anti-IgG (Biolegend, Poly4053; 1/44). Breifly, 1.5 million cells were washed twice with PBS prior to 15 min of staining with the Zombie NIR^TM^ Fixable Viability dye (BioLegend – 423105), followed by 2 washes with FACS Buffer (PBS, 5% fetal bovine serum) and fixation according to manufacturer instructions. Cells were then incubated at 4°C for 15 min with Fc Block (Rat anti-mouse CD16 and CD32; BD Pharmingen – Cat.553141) and 30 min with the antibody mixes. Both Fc Block and antibodies were diluted in the permeabilization buffer provided with the fixation kits. Finally, cells were washed twice in their respective permeabilization buffer and resuspended in FACS Buffer for acquisition on a NovoCyte Quanteon Flow Cytometer System (Agilent). The data were then analyzed on FlowJo.

### Analysis of IgA-coating of bacteria and sorting

Frozen fecal samples were homogeneized in 1 ml ice-cold PBS and centrifuged for 3 min at 100g, 4°C. Supernatant was filtered through a 70 µm straining sieve and centrifuged for 5 min at 10,000g, 4 °C. The pellet was resupended in 1 ml ice-cold PBS, the OD_600_ detected on a Nanodrop and the amount of bacteria computed as follow: 2 OD_600_ = 10^9^ bacteria. Bacteria were pelleted again for 5 min at 10,000g, 4 °C, and resuspended in 500 µl of staining buffer (PBS, 5% goat serum). After 20 min of incubation on ice, 1×10^9^ bacteria were washed with 1 ml ice-cold PBS and stained for 30 min at 4°C with 4 µg of FITC-conjugated anti-mouse IgA antibody (Southern Biotech) in 100 µl of staining buffer. Cells were then washed once and resupended in 1 ml PBS, and 100 µl of bacteria was pelleted and frozen until further analysis. Remaining bacteria where centrifuged and resuspended in 90 µl of staining buffer and 10 µL of anti-FITC MicroBeads. After 15 minutes of incubation at 4°C, bacteria were washed, resuspended in 500 µl of staining buffer and applied onto a LS column for sorting of IgA^+^ and IgA^-^ fractions with a QuadroMACS™ Separator (Miltenyi Biotec). IgA-coated and IgA-uncoated fractions were centrifuged and dry pellets were stored at -80°C until further analysis. For analysis of IgA coating of bacteria, frozen pellets were defrost on ice and washed with 1 ml of staining buffer. Since all samples were not sorted at the same time, bacteria were stained again for 30 min with 0.5 µg of FITC-conjugated anti-mouse IgA antibody (Southern Biotech) in 100 µl of staining buffer to refresh and harmonize the staining between batches. After a washing step, DNA was stained for 20 minutes with diluted 1:4000 in 200 µl of DNA staining solution (0.1 M HEPES, 0.9 % NaCl, pH 7.2). Finally, bacteria were washed twice with PBS, fixed for 20 min in 4% PFA, washed again and analyzed on a NovoCyte Quanteon Flow Cytometer System (Agilent). Data were then analyzed on FlowJo. For analysis of IgA affinity, bacteria were treated with 200 µl of PBS containing 3M NaSCN for 15 min at 4°C, 800rpm, prior to blocking and staining with FITC-conjugated anti-mouse IgA antibody as mentioned above.

### Mucus measurements

The colons were sectioned from the colon-cecum junction to the anus, and immediately fixed in freshly made Carnoy’s fixative (methanol:chloroform:glacial acetic acid, 60:30:10 v/v). The distal part of the small intestine was fixed in freshly made Carnoy’s fixative together with the half blunt end of the cecum. Fixed tissues were kept in Carnoy’s fixative for three hours and exchanged for fresh fixative for another 24 hours. The tissues were then washed in 100% methanol and kept until placed in cassettes for histology preparation. The remaining empty half of the cecum were flash frozen in liquid nitrogen and kept at -80°C until further use.

Slides were deparaffinized by submerging in xylene (Sigma-Aldrich, USA) for five minutes, followed by another xylene incubation for five minutes. Afterward, the slides were dehydrated twice in 100% ethanol for 5 minutes. The slides were then quickly washed in Milli-Q water and antigens were retrieved by submerging in antigen retrieving solution (10 mM sodium citrate, pH 6.0). The submerged sections were heated to 90°C for 10 minutes and cooldown in room temperature for 20 minutes. Slides were quickly dipped three times in Milli-Q water and blotted to remove excess liquid. To better hold liquid, a PAP pen was used to draw around the tissue area for the subsequent steps. The sections were blocked by covering the tissue in blocking buffer (1:10 goat serum (Sigma, USA) in Tris-buffered Saline (TBS; 500 mM NaCl, 50 mM Tris, pH 7.4)) and incubated for an hour at room temperature. For the primary antibody staining, the tissue was covered with a 1:200 dilution of Mucin 2 antibody (H-300) (Santa Cruz Biotechnology, USA) in blocking buffer and incubated for two hours at room temperature. Following the incubation, the slides were rinsed three times in TBS, for five minutes each. The secondary antibody staining was performed by covering the tissue with a 1:200 dilution of Alexa Fluor 488 conjugated goat anti-rabbit IgG antibody (Thermo Fisher Scientific, USA) in blocking buffer for one hour at room temperature in dark. The tissue sections were washed twice in TBS for 5 minutes, gently blotted, and covered with ProLong Gold Antifade reagent with DAPI (Invitrogen, USA), covered with cover slips and sealed with nail polish. The slides were kept at room temperature for 24 hours in dark, then kept in 4°C until imaging. The mucus layer in the sections were visualized using a Zeiss Apotome by taking pictures across fecal pellets and stitching the images together to compose a single image. Mucus layer measurements were performed by using BacSpace as described by Earle et al.

### Histological examination of intestinal tissue sections

Extent of histologic lesions was scored on a semi-quantitative basis by a trained pathologist (KE) in a blinded fashion, using a modification of the scoring system of Bugni *et al*. ^45^. Inflammation, epithelial damage, epithelial hyperplasia and dysplasia and the presence or absence of submucosa edema were scored on a scale of 1-4 according to the following table. Scores for each category were added to determine a total score. Cecum and colon were scored separately.

| Score | 0 | 1 | 2 | 3 | 4 |
| --- | --- | --- | --- | --- | --- |
| **inflammation** | Normal | Scattered PMNs or plasma cells in lamina propria or submucosa | Coalescing mucosal and/or submucosal inflammation | Widespread submucosal inflammation | Severe diffuse transmural inflammation |
| **epithelial damage** | None | Focally dilated glands and/or attenuated surface epithelium, and/or clusters of sloughed cells in lumen | Focally extensive gland dilation and/or surface epithelial attenuation, many sloughed cells in lumen | Mucosa erosions | Mucosal ulceration |
| **hyperplasia/dysplasia** | Normal | Hypertrophy and/or hyperplasia present | Cellular and/or glandular dysplasia present |  |  |
| **Submucosal edema** | Absent | Present |  |  |  |

### Mucus penetrability assay

Penetrability of the colonic mucus was assessed as described by Gustafsson et al ^21^. Briefly, colons were flushed with ice-cold oxygenated KREBS Buffer (116 mM NaCl, 1.3 mM CaCl_2_, 3.6 mM KCl, 1.4 mM KH_2_PO_4_, 23 mM NaHCO_3_, and 1.2 MgSO_4_ – Carl Roth) and opened along the mesenteric axis. The longitudinal muscle layer was removed by blunt dissection and the distal mucosa was inserted in a perfusion chamber. The basolateral chamber was filled with 0.6 µg/ml SYTO9 (Fisher Scientific - 10237582) in oxygenated KREBS Glucose Buffer (KREBS Buffer containing 10mM Glucose, 5.7 mM sodium pyruvate and 5.1 mM sodium-l-glutamate), and the apical chamber was filled with oxygenated KREBS Mannitol Buffer (KREBS Buffer, containing 10mM Mannitol, 5.7 mM sodium pyruvate and 5.1 mM sodium-l-glutamate). After 10 min incubation in the dark at room temperature, FluoSphere™ carboxylate beads (1µm, red 580/605 – Invitrogen – F882) were applied on top and let to sediment on the tissue for 5 min in the dark at room temperature. The apical chamber was then gently washed several times to remove excess of beads. The chamber was incubated for 10 min in the dark before being visualized with a microscope. For each tissue, 4-7 confocal images were taken in XY stacks from the epithelium at the bottom to the beads on top, with 5 µm-intervals between sections. Images were then analyzed with Imaris software, and the penetrability was computed by comparing the distance between the outer border of the beads and the epithelium with the distance between the most inner beads and the epithelium.

### 16S rRNA gene-based community analysis

PCR and library preparation were performed by the University of Michigan Microbiome Core lab as described by Kozich et al. ^46^. The V4 region of the 16S rRNA gene was amplified using the dual-index primers described by Kozich et al, 2013. The normalized and amplicon size evaluated samples were sequenced using an Illumina MiSeq. The raw sequences were analyzed using mothur v1.42.3 ^47^ with the included controls: PBS control during the sequencing and ZymoBIOMICS Microbial Community DNA Standard (cat# D6306) for error analysis. Sequences were aligned to the reference Silva database version 132 for contamination analysis and SPF experiments. Moreover, gnotobiotic mice sequences were also aligned to a custom reference database containing the 16S v4 region from each of the 14 bacteria members. The R package “vegan” was used to calculate the principal component analysis from the Bray-Curtis dissimilarity index.

## Notes

### Competing Interest Statement

Author M.S.D. works as a consultant and an advisory board member at Theralution GmbH, Germany.

### Summary of Updates

Reformatted for Cell press (currently under secondary review at Cell Host and Microbe December 2023).

